# Characterization of the *Chlamydomonas reinhardtii* phycosphere reveals conserved features of the plant microbiota

**DOI:** 10.1101/2021.03.04.433956

**Authors:** Paloma Durán, José Flores-Uribe, Kathrin Wippel, Pengfan Zhang, Rui Guan, Ruben Garrido-Oter

## Abstract

Microscopic algae release organic compounds to the region immediately surrounding their cells, known as the phycosphere, constituting a niche for colonization by heterotrophic bacteria. These bacteria take up algal photoassimilates and provide beneficial functions to their host, in a process that resembles the establishment of microbial communities associated with the roots and rhizospheres of land plants. Here, we characterize the microbiota of the model alga *Chlamydomonas reinhardtii* and reveal extensive taxonomic and functional overlap with the root microbiota of land plants. Reconstitution experiments using synthetic communities derived from *C. reinhardtii* and *Arabidopsis thaliana* show that phycosphere and root bacteria assemble into taxonomically equivalent communities on either host. We show that provision of diffusible metabolites is not sufficient for phycosphere community establishment, which additionally requires physical proximity to the host. Our data suggests that the microbiota of photosynthetic organisms, including green algae and flowering plants, assembles according to core ecological principles.

## Introduction

Plants associate with diverse microbes in their aerial and belowground tissues which are recruited from the surrounding environment. These microbial communities, known as the plant microbiota, provide the host with beneficial functions, such as alleviation of abiotic stresses (Xu *et al*., 2018; Berens *et al*., 2019; Simmons *et al*., 2020; Zélicourt *et al*., 2018), nutrient mobilization (Castrillo *et al*., 2017; Zhang et al., 2019; Harbort *et al*., 2020), or protection against pathogens (Durán *et al*., 2018; Carrión *et al*., 2019). Characterization of the microbiota associated with a wide range of plant species including liverworts (Alcaraz *et al*., 2018), lycopods, ferns (Yeoh *et al*., 2017), gymnosperms (Beckers *et al*., 2017; Cregger *et al*., 2018), and angiosperms (Bulgarelli *et al*., 2012; Lundberg *et al*., 2012; Edwards *et al*., 2015; Schlaeppi *et al*., 2014; Bulgarelli *et al*., 2015; Zgadzaj *et al*., 2016; Walters *et al*., 2018; Thiergart *et al*., 2020) shows a strong influence of host phylogeny as well as conserved and possibly ancestral community features. Furthermore, it has been speculated that the ability to form associations with members of these communities, such as mycorrhizal fungi, was a trait required for the colonization of land by plants 450 Mya, possibly inherited from their algal ancestor (Delaux *et al*., 2015; Knack *et al*., 2015). Algae are also known to associate with complex bacterial communities termed phycosphere microbiota, particularly in aquatic environments (Kim *et al*., 2014; Amin *et al*., 2015; Seymour *et al*., 2017; Cirri *et al*., 2019), where exchange of metabolites, including organic carbon (Moran *et al*., 2016; Wienhausen *et al*., 2017; Fu *et al*., 2020; Toyama *et al*., 2018), soluble micronutrients (Amin *et al*., 2009), vitamins (Croft *et al*., 2005; Grant *et al*., 2014; Paerl *et al*., 2017), and other molecular currencies (Teplitski *et al*., 2004; Wichard *et al*., 2015) influence algal growth and development. These parallelisms suggest that the phycosphere is analogous to the rhizosphere environment, in which secreted diffusible compounds alter soil pH, oxygen availability, concentration of antimicrobials and organic carbon, and thus support distinct microbial communities by favoring the growth of certain bacteria while restricting proliferation of others (Bell and Mitchell, 1972; Bulgarelli *et al*., 2013; Amin *et al*., 2015; Krohn-Molt *et al*., 2017; Shibl *et al*., 2020). However, it is not yet known whether the ability to assemble a complex microbiota from the surrounding soil is also conserved in soil-borne microscopic algae, and to what extent they overlap with those of vascular plants.

In this study, we characterize the microbiota of the model green alga *C. reinhardtii* (*Cr*), and show significant taxonomic and functional similarities between the root and phycosphere microbiota. In addition, we report a comprehensive, whole-genome sequenced culture collection of *Cr*-associated bacteria that includes representatives of the major taxa found in associations with land plants. We then introduce a series of gnotobiotic systems designed to reconstruct artificial phycospheres that recapitulate natural communities using synthetic communities (SynComs) assembled from bacterial isolates. Cross-inoculation and competition experiments using the model plant *Arabidopsis thaliana* (*At*) and its associated bacterial culture collection (Bai *et al*., 2015) indicate a degree of functional equivalence between phycosphere and root bacteria in associations with a photosynthetic host. Finally, we show that physical proximity between *Cr* and its microbiota is required for the establishment of fully functional phycosphere communities, suggesting that this process is not exclusively driven by the exchange of diffusible metabolites.

## Results

### *C. reinhardtii* assembles a distinct microbiota from the surrounding soil

To determine whether *Cr* shapes soil-derived bacterial communities similarly to land plants, we designed an experiment where *At* and *Cr* were grown in parallel in natural soil in the greenhouse (**Fig. S1A**). Briefly, pots containing Cologne Agricultural Soil (CAS) were inoculated with axenic *Cr* (CC1690) cultures or sowed with surface-sterilized *At* (Col-0) seeds. We then collected samples from unplanted controls and from the surface of *Cr*-inoculated pots (phycosphere fraction) at 7-day intervals, and harvested the root and rhizosphere of *At* plants after 36 days (**Methods**). Bacterial communities from all compartments were characterized by *16S* rRNA amplicon sequencing. Analysis of bacterial community profiles showed a decrease in α-diversity (Shannon index) in the phycosphere and root compartments compared to the more complex soil and rhizosphere communities (**Fig. 1A**). In addition, analysis of β-diversity revealed a significant separation by compartment, where phycosphere and root samples formed distinct clusters that were also separated from those consisting of soil and rhizosphere samples (**Fig. 1B**; 22.4% of variance; *P*<0.001). Further inspection of amplicon profiles showed an overlap between root- and phycosphere-associated communities along the second and third components (**Fig. 1C**), suggesting similarities between the bacterial communities that associate with *Cr* phycospheres and *At* roots.

**Figure 1.**
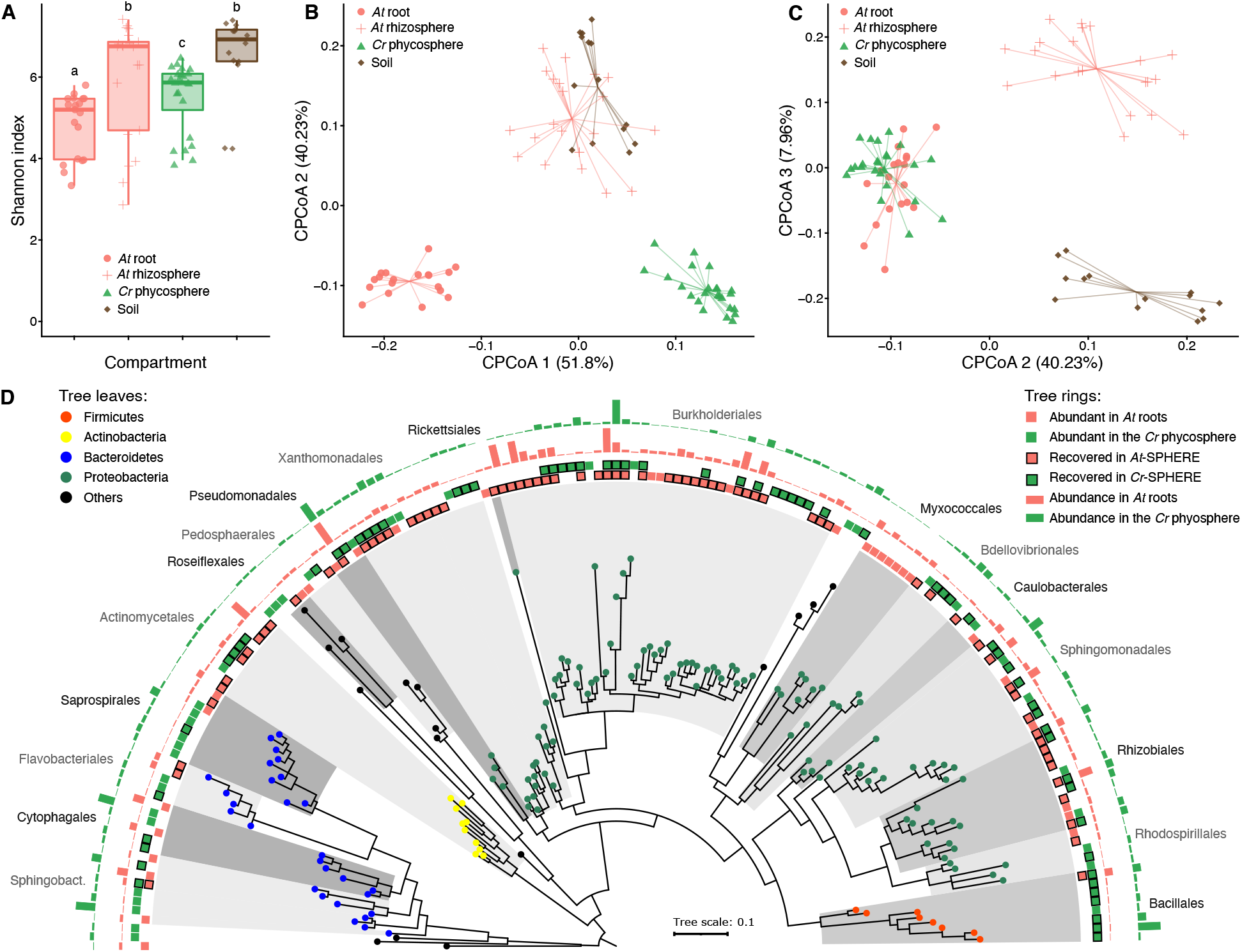
Comparison of bacterial community structures associated with *At* roots and the *Cr* phycosphere in natural soil. (**A**) Alpha diversity estimates of soil, rhizosphere, root and phycosphere samples from *At* and *Cr* grown in CAS soil in the greenhouse. (**B-C**) PCoA of Bray-Curtis dissimilarities constrained by compartment (22.4% of variance explained; *P*<0.001). A separation between root, phycosphere and soil-derived samples can be observed in the first two components (**B**), while the root and phycosphere communities cluster together in the second and third PCoA axes (**C**). (**D**) Phylogeny of *16S* rRNA sequences of the most abundant OTUs found in *At* roots and *Cr* phycosphere community profiles. Leaf nodes are colored by taxonomic affiliation (phylum level). The two innermost rings (colored squares) represent abundant OTUs in each compartment. Squares highlighted with a black contour correspond to OTUs for which at least one representative bacterial strain exists in the IRL or IPL culture collections. The two outermost rings (barplots) represent log-transformed relative abundances of each OTU in *At* root or *Cr* phycosphere samples.

To characterize the dynamics of these microbiota assembly processes, we analyzed the time-series data from soil and phycosphere and end-point community profiles from *At* roots. This revealed a gradual recruitment of bacterial taxa from soil, leading to the formation of distinct phycosphere communities that become significantly differentiated 21 days after inoculation, which is of comparable to that observed in *At* root-associated communities at day 36 (**Fig. S2A**). Subsequent enrichment analysis of amplicon sequence variants (ASVs) in each compartment, compared to unplanted soil, showed an increase in the relative abundance of *Cr*- and *At*-enriched ASVs in phycosphere and root samples, respectively. In contrast, total relative abundance of soil-enriched ASVs progressively decreased in host-associated compartments, while remaining stable in unplanted soil (**Fig. S2B-D**). Although the magnitude of the changes in bacterial community composition in the phycosphere diminishes over time, it remains unclear whether these communities reach a steady state over the duration of the experiment. Taken together, these results indicate that, similarly to *At, Cr* is able to recruit a subset of bacterial taxa from the surrounding soil and assemble a distinct microbiota.

### The *C. reinhardtii* phycosphere and the plant root share a core microbiota

Given the observed similarities between phycosphere and root communities (**Fig. 1C**), we compared the most abundant taxonomic groups found in association with the two photosynthetic hosts. We found a significant overlap between Operational Taxonomic Units (OTUs) with the highest relative abundances in either phycosphere or root samples (**Fig. 1D**; >0.1% relative abundance; 32% shared; *P*<0.001), which included members of every bacterial order except Myxococcales, which were only found in large relative abundances in *At* root samples (**Supplementary Data 1**). In line with previous descriptions of the *At* root microbiota, we observed that these host-associated communities were dominated by Proteobacteria, and also included members of the Actinobacteria, Bacteroidetes, and Firmicutes phyla. At this taxonomic level, the major difference between the two photosynthetic hosts was given by a lower contribution of Actinobacteria and a larger relative abundance of Firmicutes in the *Cr* phycosphere compared to the *At* root compartment (**Fig. 1D**). Given that this latter phylum is most abundant in soil, this difference may be due to the difficulty of fully separating soil particles from the phycosphere fraction during sample collection.

Next, we sought to assess whether the observed overlap in community structures between *Cr* and *At* could be extended to other land plant lineages. We performed a meta-analysis, broadening our study to include samples from phylogenetically diverse plant species found in a natural site, including lycopods, ferns, gymnosperms, and angiosperms (Yeoh *et al*., 2017), as well as the model legume *Lotus japonicus* (*Lj*) grown in CAS soil in the greenhouse (Thiergart *et al*., 2019; Harbort *et al*., 2020). First, we determined which taxonomic groups were present in each plant species (≥80% occupancy and ≥0.1% average relative abundance) and identified a total of six bacterial orders that consistently colonize plant roots (i.e., found in every host species). These taxa include Caulobacterales, Rhizobiales, Sphingomonadales, Burkholderiales, Xanthomonadales (Proteobacteria), and Chitinophagales (Bacteroidetes). We observed that the aggregated relative abundance of these six bacterial orders accounted for 39% of their respective communities on average (**Fig. 2**). Interestingly, these taxa were also found among the most abundant in the *Cr* phycosphere (45% aggregated relative abundance), indicating that they are also able to associate with *Cr*. These results suggest the existence of a common principle for microbiota assembly across a wide phylogenetic range of photosynthetic hosts, which includes uni- and multicellular eukaryotic organisms.

**Figure 2.**
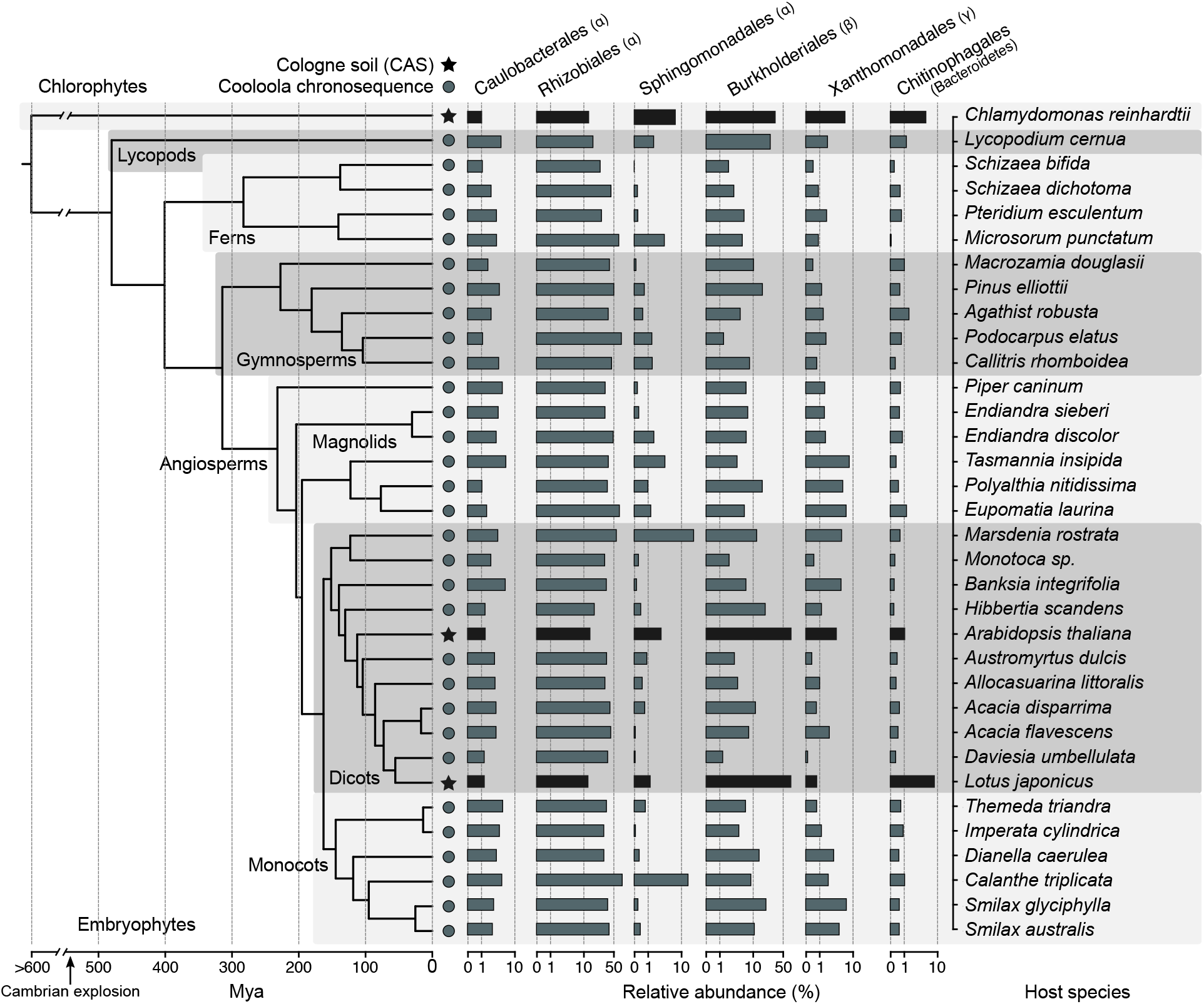
Conservation of bacterial orders of the root and phycosphere microbiota across photosynthetic organisms. Phylogeny inferred from a multiple sequence alignment of the ribulose-bisphosphate carboxylase gene (*rbcL*) of 35 plant species and *Chlamydomonas reinhardtii*. The barplots represent the average aggregated relative abundance of the six bacterial orders found to be present in the root microbiota of each plant species (80% occupancy and ≥0.1% average relative abundance). Leaf nodes depicted with a star symbol denote community profiles of plants grown in CAS soil in the greenhouse (Thiergart *et al*., 2019; Harbort *et al*., 2020), whereas those marked with a circle were obtained from plants sampled at the Cooloola natural site chronosequence (Yeoh *et al*., 2017).

### Reconstitution of phycosphere communities using reductionist approaches

After the characterization of phycosphere-associated bacterial communities in natural soil, we sought to develop systems of reduced complexity that would allow controlled perturbation of environmental parameters, and targeted manipulation of microbial community composition. First, we established a mesocosm system using soil-derived microbial communities as start inocula (**Fig. S1B**). We co-inoculated axenic *Cr* (CC1690) cultures with microbial extracts from two soil types (CAS and Golm) in two different carbon-free media (TP and B&D), which ensures that the only source of organic carbon to sustain bacterial growth is derived from *Cr* photosynthetic activity (**Methods**). These phycosphere mesocosms were then incubated under continuous light for 11 days, during which we assessed *Cr* growth using cell counts, and profiled bacterial communities *via 16S* rRNA amplicon sequencing. In this system*, Cr* was able to steadily grow without a detrimental impact from co-inoculation with soil-derived bacterial extracts (**Fig. S3A**). Analysis of diversity showed that *Cr* was able to shape soil-derived bacterial communities within the first 4 days, compared to the starting inocula, and that these phycosphere communities remained stable until the end of the experiment (**Fig. 3**). Interestingly, cultivation of soil-derived bacteria in the absence of organic carbon or supplemented with Artificial Photosynthates (AP; **Methods**) led to significantly differentiated bacterial communities (**Fig. 3A**; 17.9% of variance; *P*<0.001). In addition, inoculation of soil-derived bacteria with heat-killed *Cr* cultures was not sufficient to recapitulate this community shift (**Fig. S3B**), suggesting that the presence of live and metabolically active *Cr* is required for the establishment of synthetic phycospheres. We then tested whether larger eukaryotic microorganisms present in the soil microbial extracts, such as other unicellular algae or fungi, were also contributing to the observed changes in bacterial composition. A separate experiment, where microbial inocula were filtered through a 5 μm pore-size membrane, showed similar bacterial community shifts compared to non-filtered extracts (**Fig. S3C**). Similar to the results obtained using natural soil, the aggregated relative abundance of *Cr*-associated ASVs in the synthetic phycosphere samples increased over time, whereas ASVs enriched in the bacteria only control samples consistently decreased (**Fig. 3B**). At the end of the experiment (day 11), the relative abundance of *Cr*-enriched ASVs accounted for 94% of the entire phycosphere community, in contrast to a lower contribution observed in the natural soil system (**Fig. S2B**; 60% relative abundance at day 36). This pattern could be a consequence of the unintended depletion of bacteria that are not capable of metabolizing *Cr*-secreted photoassimilates in a liquid environment, and in these specific culture media. Finally, an independent mesocosm experiment using day/night light cycles showed delayed but similar patterns to those using continuous light, indicating that phycosphere community establishment may be independent of *Cr* culture synchronization (**Fig. S3D**).

**Figure 3.**
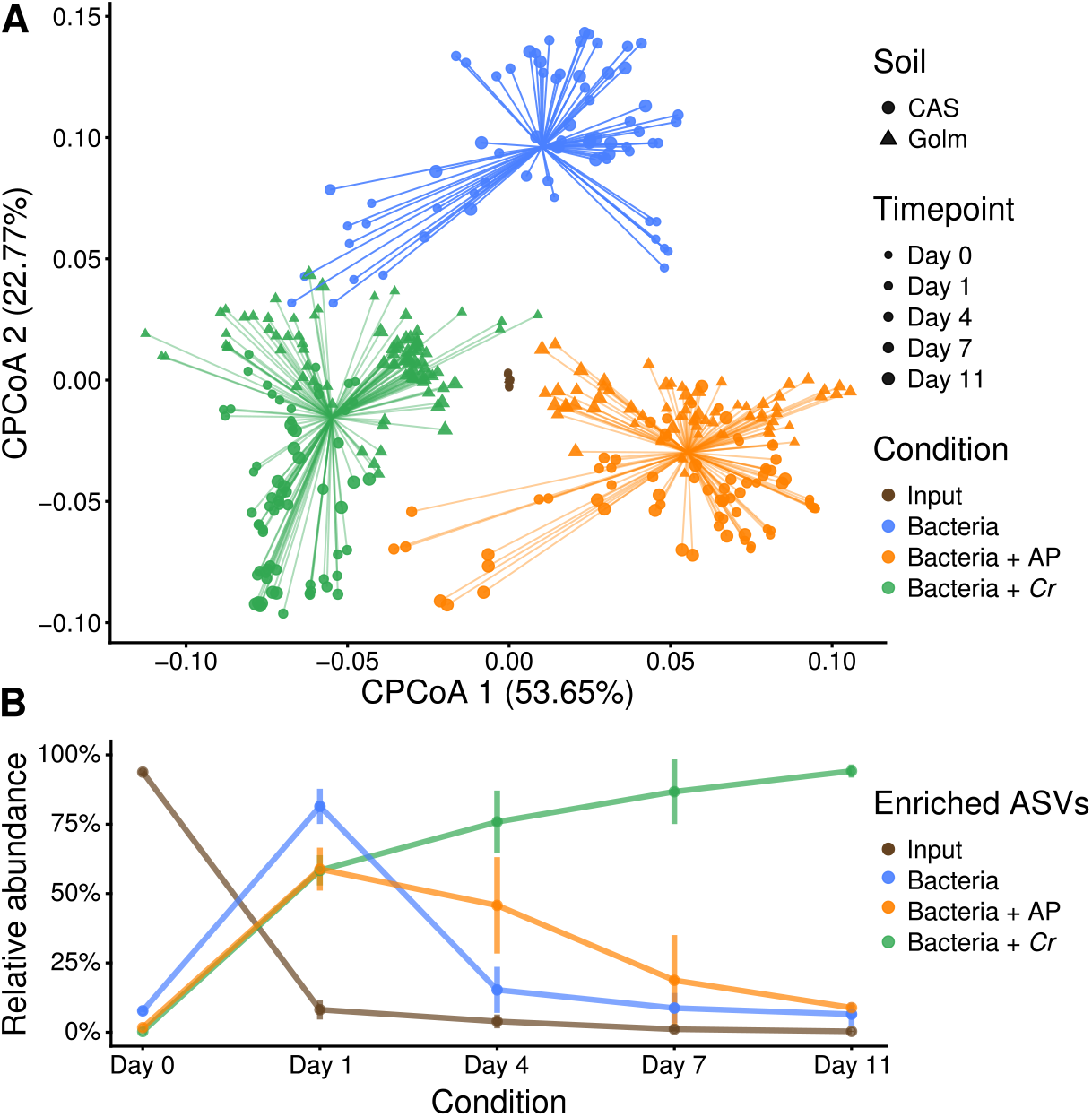
Mesocosm experiments recapitulate the establishment of phycosphere communities by *Cr* across soil types and growth media. (**A**) PCoA analysis of Bray-Curtis dissimilarities constrained by condition (17.9% of variance; *P*<0.001) show a significant separation between start inocula (soil washes, depicted in brown), phycosphere communities (green), and soil washes incubated in minimal media (blue), or media supplemented with artificial photoassimilates (APs, depicted in orange). (**B**) Dynamic changes in the phycosphere community composition in terms of the aggregate relative abundances of ASVs enriched in each condition with respect to the start inocula.

Next, we aimed to control community composition in this reductionist system by establishing a *Cr*-associated bacterial culture collection following a similar approach as reported in previous studies with land plants (Bai *et al*., 2015; Lebeis *et al*., 2015; Eida *et al*., 2018; Garrido-Oter *et al*., 2018; Wippel *et al*., 2021; Zhang *et al*., 2021a). We employed a limiting dilution approach using 7 day-old *Cr* phycospheres derived from CAS soil bacteria incubated in two minimal media (TP and B&D; **Methods**). The resulting sequence-indexed phycosphere bacterial library (*Cr*-IPL) contained a total of 1,645 colony forming units (CFUs), which were taxonomically characterized by *16S* rRNA amplicon sequencing. Comparison of these sequencing data with the community profiling of soil phycospheres revealed that we were able to recover 62% of the most abundant bacterial OTUs found in natural communities (**Fig. S4A**; **Supplementary Data 2**). Recovered OTUs accounted for up to 63% of the cumulative relative abundance of the entire culture-independent community, indicating that our collection is taxonomically representative of *Cr* phycosphere microbiota. These results are comparable to the recovery rates observed in previously reported culture collections from different plant species (e.g., 57% for *A. thaliana*, Bai *et al*., 2015; 69% for rice, Zhang *et al*., 2019; 53% for *L. japonicus;* Wippel *et al*., 2021).

To establish a core collection of phycosphere bacteria, we selected a taxonomically representative set of strains from the *Cr*-IPL covering all major taxonomic groups found in the culture-independent community profiles and subjected them to whole-genome sequencing (**Methods**). In total, we sequenced the genomes of 185 bacterial isolates, classified into 42 phylogroups (97% average nucleotide identity), belonging to 5 phyla and 15 families (**Supplementary Data 3**). Next, we performed comparative analyses of the genomes from the phycosphere core collection (*Cr*-SPHERE) with those established from soil, roots of *A. thaliana*, and roots and nodules of *L. japonicus* (*At*- and *Lj*-SPHERE) grown in the same soil (CAS). A whole-genome phylogeny of these bacterial strains showed that all major taxonomic groups that included root-derived isolates were also represented in the *Cr*-SPHERE collection, but not in the soil collection (**Fig. S4B**). Importantly, the phycosphere collection also included multiple representatives of each of the six bacterial orders that were found to consistently colonize plant roots in natural environments (**Fig. 2**). Next, we assessed the functional potential encoded in the genomes of the sequenced phycosphere bacteria using the KEGG orthology database as a reference (Kanehisa *et al*., 2014). Principal coordinates analysis (PCoA) of functional distances showed that bacterial taxonomy accounted for most of the variance of the data (58.63%; *P*<0.001), compared to a much smaller impact of the host of origin of the genomes (4.22% of variance; *P*<0.001; **Fig S4C**).

Next, we tested whether synthetic communities formed by isolates from the *Cr*-SPHERE collection could recapitulate assembly patterns of natural phycospheres under laboratory conditions. Axenic *Cr* cultures (CC1690) were inoculated with a bacterial SynCom composed of 26 strains that could be distinguished at the *16S* level and contained representative members of all major phycosphere taxonomic groups (**Fig. S1D**; **Supplementary Data 4**). Assessment of *Cr* growth using chlorophyll fluorescence and cell counts showed that the presence of the bacterial SynCom had no consistent beneficial or detrimental impact on *Cr* proliferation in this system (**Fig. 4A-B**), similarly to what we observed in mesocosms (**Fig. S3A**). Analysis of time-course amplicon profiles showed that *Cr* assembled a characteristic phycosphere community within the first 4 days of co-inoculation, which was significantly separated from both, start inocula and bacterial SynComs alone (**Fig. 4C-D**). Together, these results demonstrate that we can recapitulate *Cr* assembly of distinct phycosphere communities in natural soils using culture-dependent and -independent gnotobiotic systems.

**Figure 4.**
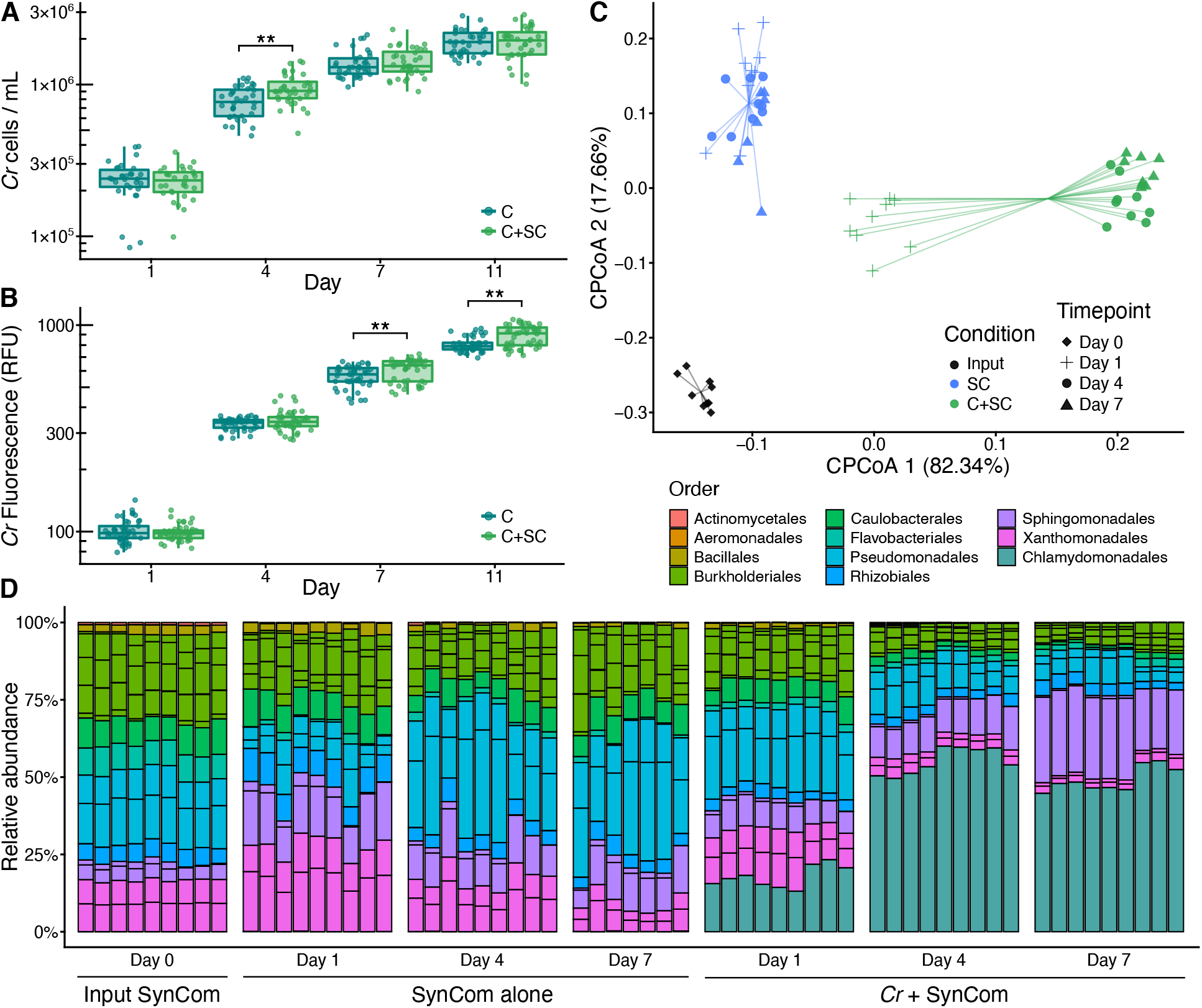
Phycosphere reconstitution using bacterial SynComs derived from the *Cr*-SPHERE core culture collection. (**A**) Beta-diversity analysis (CPCoA of Bray-Curtis dissimilarities; 40.4% of the variance; P<0.001) of samples obtained from a liquid-based gnotobiotic system, showing a significant separation between input SynCom samples (black), synthetic phycospheres (light green), and SynCom only controls (blue). (**B**) Bar charts showing relative abundances of individual SynCom members across conditions and timepoints (colored by their taxonomic affiliation at the order level). (**C-D**) *Chlamydomonas* growth in the gnotobiotic system axenically (dark green) or in co-inoculation with the bacterial SynCom (light green), measured as algal cell densities (**C**), and relative chlorophyll fluorescence (**D**).

### *Cr*- and *At*-derived SynComs form taxonomically equivalent communities on either host

Given the similarity between phycosphere and root communities observed in natural soils (**Fig. 1**), and the taxonomic and functional overlap across genomes from their corresponding core collections (**Fig. S4B-C**), we hypothesized that SynComs with the same taxonomic composition would assemble into similar communities, regardless of their origin. To test this hypothesis, we used a soil-based gnotobiotic system in which we could grow *Cr* and *At* in parallel, in addition to the previously described liquid-based system (**Methods**). We designed taxonomically-paired SynComs composed of strains from either the IPL (*Cr*-SPHERE) or IRL (*At*-SPHERE) bacterial culture collections. In these SynComs we included one representative strain from each bacterial family shared between the two collections (*n*=9), ensuring that they could be differentiated by their *16S* rRNA sequences (**Supplementary Data 4**). We then inoculated axenic *Cr* cultures and *At* seeds with either IPL, IRL or mixed (IPL+IRL) SynComs and allowed to colonize either host for four weeks (**Fig. S1E**). Next, we harvested the root, soil, and phycosphere fractions, measured host growth, and performed *16S* rRNA amplicon sequencing (**Methods**). Assessment of growth parameters (cell counts for bacteria and *Cr*, chlorophyll content for *Cr* and shoot fresh weight for *At*) showed no significant differences across SynCom treatments (**Fig. S5**). However, analysis of community profiles of the mixed SynComs showed that *Cr* and *At* assemble distinct communities that could also be clearly separated from unplanted soil (**Fig. 5A**). Similar to what we observed in natural soil (**Fig. 1C**), there was an overlap between phycosphere and root samples, which clustered together along the second and third components (**Fig. 5B**). Interestingly, analysis of community composition at the family level showed that all SynComs (*Cr*-, *At*-derived, and mixed) formed taxonomically indistinguishable root or phycosphere communities, independently of their host of origin (**Fig. 5A-B**). Furthermore, analysis of aggregated relative abundances from mixed communities showed that phycosphere-derived strains could successfully colonize *At* roots (48.32% relative abundance), and root-derived strains established associations with *Cr* in both soil and liquid systems (42.94% and 25.70% relative abundance, respectively; **Fig. 5C-D**). Despite this capacity for ectopic colonization, we observed significant signatures of host preference in SynComs from the two culture collections, indicated by the fact that *Cr*-derived strains reached higher aggregated relative abundances in the phycosphere compared to the root, while the opposite pattern was identified for *At*-derived bacteria (**Fig. 5C**). This tendency was accentuated in the liquid system, where *Cr* bacteria outcompeted *At* strains in the presence of the algae but not when they were incubated alone (**Fig. 5D**). Taken together, these results suggest the presence of conserved features in bacterial members of the *Cr* and *At* microbiota at a high taxonomic level, with signatures of host preference at the strain level.

**Figure 5.**
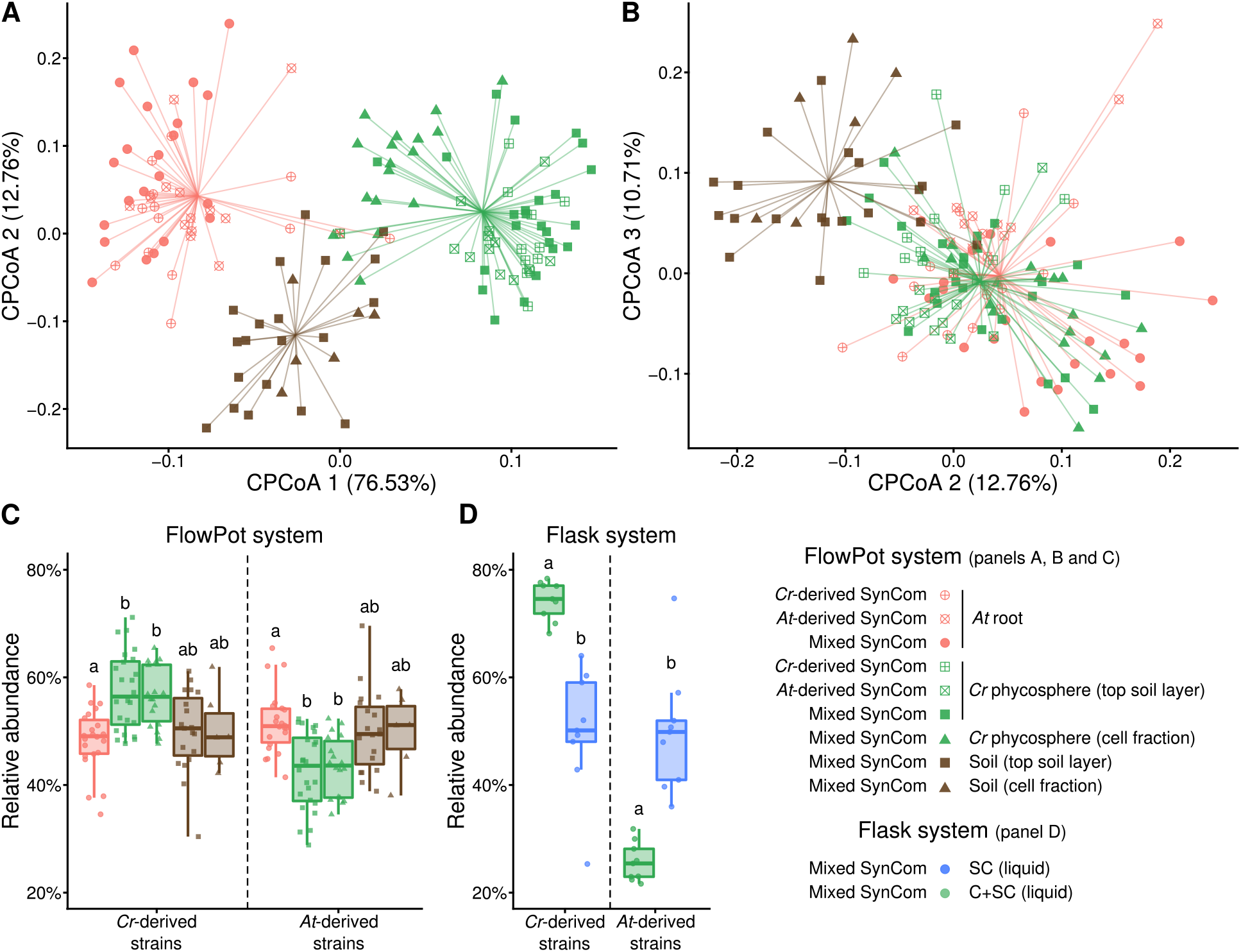
Root and phycosphere bacteria colonize *At* and *Cr* and assemble into taxonomically equivalent communities. (**A-B**) Beta diversity analysis of soil, root, and phycosphere community profiles obtained from gnotobiotic *At* and *Cr*, inoculated with bacterial SynComs derived from *At* roots (*At*-SPHERE), *Cr* (*Cr*-SPHERE) or mixed (*At*- and *Cr*-SPHERE), grown in the FlowPot system analysis (CPCoA of Bray-Curtis dissimilarities aggregated at the family level; 16.4% of the variance; *P*<0.001). Similar as in natural soils (**Fig. 1B-C**), root and phycosphere samples were significantly separated from soil and from each other in the first two axes, while overlapping in the second and third components. (**C-D**) Aggregated relative abundances of *At*- and *Cr*-derived strains in the mixed SynCom show ectopic colonization and signatures of host preference in a soil-derived (FlowPot, panel **C**), and liquid-based (flask, panel **D**) gnotobiotic system.

### Physical proximity is required for the assembly of phycosphere communities and promotion of *Cr* growth

Next, we sought to investigate whether the observed formation of distinct phycosphere communities (**Figs. 1**, **3** and **4**) is driven by the secretion of diffusible photoassimilates and to what extent physical proximity to bacteria is required to establish other forms of interactions. To test this hypothesis, we developed a gnotobiotic split co-cultivation system where synthetic phycospheres could be grown photoautotrophically (**Fig. S1F**). In this system, two growth chambers were connected through a 0.22 μm-pore polyvinylidene fluoride (PVDF) membrane that allows diffusion of compounds but not passage of bacterial or algal cells (**Methods**). We co-cultivated axenic *Cr* cultures (C), bacterial SynComs (SC), and synthetic phycospheres (C+SC) in these split chambers containing minimal carbon-free media (TP) in multiple pair-wise combinations (**Fig. S1F**; **Supplementary Data 4**). Analysis of *16S* rRNA amplicon profiles after 7 days of incubation revealed that SC and C+SC samples were distinguishable from the input bacterial SynComs (**Fig. 6A**). In addition, samples clustered according to the presence of *Cr* in the same compartment, causing SC and C+SC samples to be significantly separated, independently of the community present in the neighboring chamber (**Fig. 6A**, indicated by colors; 21.4% of variance; *P*<0.001). Comparison of amplicon profiles of samples taken from chambers containing C+SC further showed a significant impact of the content of the neighboring compartment in community structures (**Fig. 6B**, indicated by shapes; 39.5% of variance; *P*<0.001). Interestingly, we also observed that the presence of *Cr* in the neighboring compartment was sufficient to change SC communities where the bacterial SynCom was incubated alone (**Fig. 6C**; **SC**|C or **SC**|C+SC *versus* **SC**|–; *P*=0.001), possibly by secreting diffusible compounds or inducing changes in the composition of the culture medium (e.g., minerals, pH). Furthermore, SC communities where *Cr* was present in the neighboring compartment could be differentiated depending on whether *Cr* was in direct contact with bacteria or grown axenically (**Fig. 6C**; **SC**|C *versus* **SC**|C+SC). These community shifts could be explained by competition for diffusible metabolites with the neighboring compartment containing the SynCom together with the algae (C+SC), or by physiological changes in *Cr* induced by physical proximity with bacteria.

**Figure 6.**
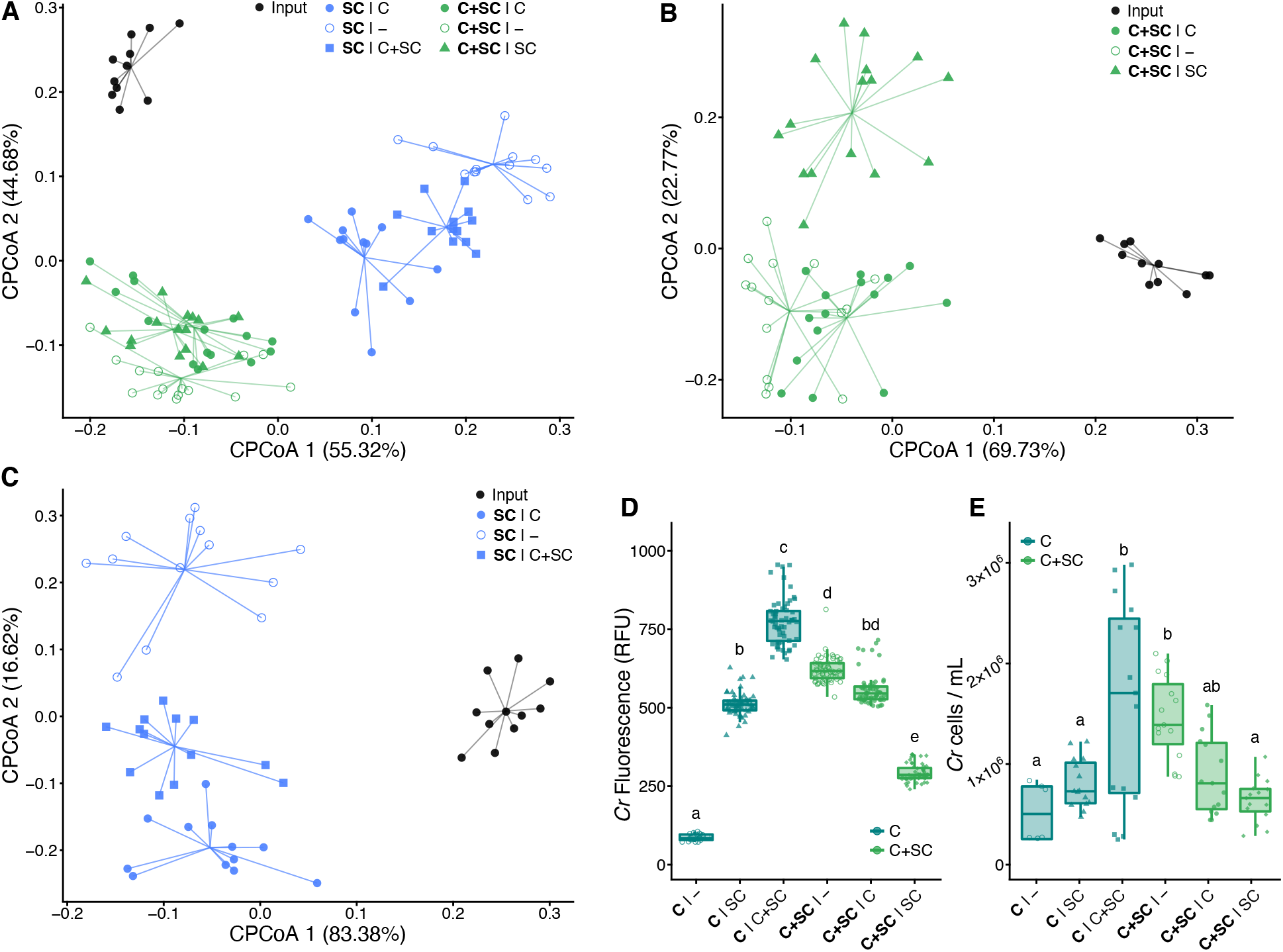
Physical proximity to *Cr* is required for the establishment of phycosphere bacterial communities. Beta-diversity analyses of Bray-Curtis dissimilarities of SynComs grown in a split gnotobiotic system show a significant separation of samples according physical proximity to *Cr* (21% of variance; *P*<0.001, **A**), or the content of the neighboring vessel (39.4-39.5% of variance; *P*<0.001, panels **B-C**). (**D-E**) *Cr* growth across conditions measured using relative chlorophyll fluorescence (RFU; panel **D**) and algal cell densities (panel **E**).

In parallel to bacterial community profiles, we assessed *Cr* growth by measuring chlorophyll fluorescence and algal cell counts in all vessels (**Methods**). We observed significant differences in the growth of axenic *Cr* cultures depending on the contents of the neighboring chamber, where the bacterial SynCom alone (**C**|SC) had a positive impact on the microalgae compared to the control (**C**|–; **Fig. 6D-E**). Remarkably, the presence of a synthetic phycosphere in the neighboring compartment had the strongest positive impact on axenic *Cr* cultures (**C**|C+SC; **Fig. 6D-E**), suggesting that changes in bacterial community composition driven by physical proximity to *Cr* lead to a beneficial impact on algal growth. In addition, chlorophyll fluorescence and cell counts of synthetic phycospheres (C+SC) were higher when no other microorganisms were incubated in the neighboring chamber (**C+SC**|– *versus* **C+SC**|C or **C+SC**|SC; **Fig. 6D-E**), possibly due to competition for diffusible compounds. An additional full-factorial replicate experiment using a modified version of this split co-cultivation system (**Methods**) showed consistent results both in community structures and *Cr* growth parameters (**Fig. S6**), despite of a large technical variation in cell density measurements (**Fig. 6D**). Together, these results indicate that physical proximity of bacteria to *Cr* is required for assembly and growth of phycosphere communities, which in turn may benefit host growth by providing metabolites and / or other compounds including carbon dioxide, which in this experimental setup is likely limiting autotrophic growth of *Cr*. Future experimentation with synthetic phycospheres composed by SynComs designed using combinatorial approaches, coupled with metabolomic and transcriptomic profiling, will be needed to decipher the molecular and genetic mechanisms driving these interactions.

## Discussion

Microscopic algae release photoassimilated carbon to the diffusible layer immediately surrounding their cells, which constitutes a niche for heterotrophic bacteria. Microbes from the surrounding environment compete for colonization of this niche and assemble into complex communities that play important roles in global carbon and nutrient fluxes. These ecological interactions have been well studied in aquatic environments, where each year approximately 20 Gt of organic carbon fixed by phytoplankton are taken up by heterotrophic bacteria (Moran *et al*., 2016), which can account for up to 82% of all algal-derived organic matter (Horňák *et al*., 2017). For multiple species of green algae, optimal growth in turn requires interactions with their associated phycosphere bacteria, which can provide beneficial services to their host, such as mobilization of non-soluble iron (Amin *et al*., 2009), or exogenous biosynthesis of organic compounds such as vitamins (Croft *et al*., 2005; Paerl *et al*., 2017). Despite the known importance of these interactions in marine environments, the role of algae-bacterial associations in terrestrial ecosystems remains understudied. This gap in our understanding could be explained by the fact that aquatic phytoplankton are more readily noticed and more amenable to systematic study compared to edaphic microalgae. However, exploring the role of soil-borne unicellular photosynthetic organisms as hosts of complex microbial communities could expand our understanding of carbon and energy fluxes in terrestrial ecosystems.

The results from our culture-independent and gnotobiotic experiments using the ubiquitous algae *Cr*, which was originally isolated from soil (Sasso *et al*., 2018), illustrate that green algae can recruit and sustain the growth of heterotrophic, soil-borne bacteria. This process resembles the establishment of the microbial communities that associate with the roots and rhizospheres of land plants, suggesting common organizational principles shared between chlorophytes and embryophytes. Our in-depth characterization of the *Cr* microbiota shows clear differences as well as striking similarities in the taxonomic affiliation of abundant root and phycosphere community members (**Fig. 1D**). Notably, these similarities are found despite biochemical differences between extracellular organic carbon compounds released by *At* roots and *Cr*, as well as by differences in cell wall composition, which in the case of the plant root mostly consists of complex polysaccharides such as cellulose, whereas in *Cr* it is primarily composed of (glyco)proteins (Harris, 2009). Among the bacterial lineages shared between the root and phycosphere microbiota, we found groups that are known to establish intimate interactions with multicellular plants, ranging from symbiotic to pathogenic, such as Rhizobia, Pseudomonas, Burkholderia, or Xanthomonas (Suarez-Moreno, 2014; Garrido-Oter *et al*., 2018; Karasov *et al*., 2018; Timilsina *et al*., 2020). Meta-analyses of available data from multiple studies further confirm this pattern by revealing the presence of a set of six bacterial orders, found as abundant members not only in the root communities of all analyzed land plants, but also in the *Cr* phycosphere (**Fig. 2**). These findings suggest that the capacity to associate with a wide range of photosynthetic organisms is a common trait of these core bacterial taxa, which might predate the emergence of more specialized forms of interaction with their host. This hypothesis was implicitly tested in our cross-inoculation gnotobiotic experiments, where bacterial strains originally isolated from the roots of *At* or the phycosphere of *Cr* competed for colonization of either host (**Fig. S1E**). The observation that *Cr*-derived strains could colonize *At* roots in a competition setup, whereas *At*-derived bacterial SynComs also populated *Cr* phycospheres (**Fig. 5C**) supports the existence of shared bacterial traits for establishing general associations with photosynthetic hosts. Despite these patterns of ectopic colonization, we also detected significant signatures of host preference, illustrated by the observation that native bacterial SynComs outcompeted non-native strains in the presence of either host, but not in their absence (**Fig. 5C**). These findings are in line with a recent comparative microbiota study where similar results were observed for bacterial commensals from two species of land plants (*A. thaliana* and *L. japonicus;* Wippel *et al*., 2021). In addition, SynComs composed of strains exclusively derived from the *At-* or the *Cr*-SPHERE collections, assembled into taxonomically equivalent communities on either host, which were indistinguishable at the family level (**Fig. 5A-B**). Together, our findings suggest that these bacterial taxa have in common the ability to assemble into robust communities and associate with a wide range of photosynthetic organisms, including unicellular algae and flowering plants.

Carbon is assumed to be the main factor limiting bacterial growth in soil (Demoling *et al*., 2007). Thus, secretion of organic carbon compounds by photosynthetic organisms constitutes a strong cue for the assembly of soil-derived microbial communities (Bulgarelli *et al*., 2013; Zhalnina *et al*., 2018; Huang *et al*., 2019). The observed similarities between the root and phycosphere microbiota at a high taxonomic level suggest that the release of photoassimilates acts as a first organizing principle driving the formation of these communities. This hypothesis is also supported by a recent study with marine bacterial mesocosms where community composition could be partially predicted by the addition of phytoplankton metabolites (Fu *et al*., 2020). However, the results from our split system (**Fig. S1F** and **Fig. 6**), where bacterial SynComs formed distinct communities and had a beneficial effect on *Cr* growth depending on their physical proximity, indicate that the provision of diffusible carbon compounds is not sufficient to explain the observed patterns of microbial diversity. In addition, shed *Cr* cell wall components, which may not be diffusible through the 0.22 μm-pore membrane, could be degraded by bacteria only in close proximity. The importance of proximity to the algal cells could also be a consequence of gradients in concentrations and variations in the diffusivity of different compounds, which in aquatic environments is predicted to cause highly chemotactic, copiotrophic bacterial populations to outcompete low-motility oligotrophic ones (Smriga *et al*., 2016). Together with the algal growth data, the observed variations in SynCom structures suggest that, in addition to physical proximity, bi-directional exchange of metabolic currencies and / or molecular signals may be required for the assembly and sustained growth of a phycosphere microbiota capable of providing beneficial functions to their host. Future experimentation using this system will be aimed at elucidating core molecular and ecological principles that govern interactions between photosynthetic organisms and their microbiota.

## Methods

### *Cr* culture conditions

*Cr* CC1690 cells were grown photoautotrophically in TP (Kropat *et al*., 2011), TP10 or B&D medium (Broughton and Dilworth, 1971) at 25 °C, and an illumination of 125 μmol m-2 s-1 under continuous light conditions. Cultures were kept in a rotatory shaker at 70 RPM. Cells in the mid-logarithmic phase were used as inocula for the different experiments. Cell growth was determined either by measuring samples in a Multisizer 4e Coulter counter (Beckman Coulter Inc., California, USA) or using an Infinite M200Pro (TECAN Austria GmbH, Grödig, Austria) plate reader to determine either absorbance at 750 nm or chlorophyll fluorescence (excitation 440/9 nm, emission 680/20 nm).

### Greenhouse experiment

*Arabidopsis thaliana* Col-0 seeds were surface-sterilized in 70% ethanol for 10 min followed by a brief wash with 100% ethanol (1 min), a wash with 3% NaClO (1 min) and five subsequent washes with sterile water. Seeds imbibed in sterile water were stratified for four days at 4°C in the dark. Five seeds were then directly sown onto the surface of pots containing Cologne Agricultural Soil (CAS) by pipetting one seed at a time.

After 36 days plants were harvested similarly to previously reported protocols (Thiergart *et al*., 2020). Briefly, and plant roots were manually separated from the surrounding soil, until only tightly adhered soil particles were left. Then, roots were separated from their shoot and placed in a Falcon tube with 10 mL of deionized sterile water. After ten inversions, the roots were transferred to another Falcon tube and further processed, while leftover wash-off was centrifuged at 4,000×g for 10 min. The supernatant was discarded and the pellet was resuspended and transferred to a new 2-mL screw-cap tube. This tube was centrifuged at 20,000 RPM for 10 min, the supernatant was discarded and the pellet snap-frozen in liquid nitrogen and stored for further processing (rhizosphere compartment). Root systems were then washed successively in 80% EtOH and 3% NaOCl to further clean the root surfaces from living microorganisms and subsequently washed three times (1 min each) in sterile water. These microbe-enriched root fractions were transferred to 2-mL screw-cap tubes for further processing (**Fig S1A**).

Liquid TP cultures of 7-day old *Cr* (CC1690) were washed by sequential centrifugation at 5,000×g for 5 min and resuspended in 50 mL of MgCl_2_, to an average 1.4×10^6^ cells/mL across biological replicates, to be used as inocula for CAS pots. Samples from the surface of the *Cr*-inoculated pots were collected using an ethanol-washed metal spatula at 7, 14, 21, 28, and 36 days post-inoculation. Unplanted pots containing CAS were used to collect surface samples as mock-treatment control right after inoculation (day 0) and at the same time points as *Cr*-inoculated pots. The position of the pots in the trays was shuffled periodically to minimize edge and location effects (**Fig S1A**). Sterile petri dishes were placed at the bottom of each pot, which were then watered from the top at inoculation time with 50 mL of MgCl_2_, and then by adding sterile MilliQ water every 2-3 days in the petri dishes, and kept in the greenhouse under long-day conditions (16/8 h light/dark). Collected samples were snap-frozen using liquid nitrogen and stored at −80 °C until further processing.

### Microbial soil wash preparation

Soil samples (5 g) from CAS or GOLM soil were collected in Falcon tubes and manually resuspended onto 30 mL of sterile 1x Tris-EDTA (TE) supplemented with 0.1% of Triton X-100 (SERVA Electrophoresis GmbH, Heidelberg, Germany). The solution was then homogenized by inversion at 40 RPM for 30 min in a rotary mixer and centrifuged for 1 min at 1500 RPM to remove bigger soil particles. Afterwards, the supernatant was transferred to a new Falcon tube and centrifuged at 4,000×g for 20 min. After centrifugation the supernatant was discarded and the pellet resuspended in 50 mL of the final medium. Cell concentration was then determined using either a hemocytometer or the Multisizer 4e.

*Cr* cells from an axenic culture were inoculated to a density of 10^5^ cells/mL into 50 mL of TP or B&D medium in triplicate in 200mL flasks. An estimate of 10^9^ cells from the microbial soil wash were added to the same flasks and incubated for 11 days as described above. Controls consisted in flasks, wrapped in aluminum foil to prevent the pass of light, containing the same growth media as the one used for the *Cr* cultures with and without artificial photosynthates (AP; Baudoin *et al*., 2003). Samples were collected for DNA extraction and cell counts determination at 0, 1, 4, 7, and 11 days post inoculation (**Fig S1B**). These experiments were repeated in three biologically independent experiments, per soil type and growth media.

### DNA extraction from soil samples

Total DNA was extracted from the aforementioned samples using the FastDNA™ SPIN Kit for Soil following instructions from the manufacturer (MP Biomedicals, Solon, USA). DNA samples were eluted in 50 μL nuclease-free water and used for microbial community profiling.

### DNA extraction from liquid samples

DNA from liquid samples was extracted using alkaline lysis (Bai *et al*., 2015). Briefly, 12 μL of the sample were diluted in 20 μL of Buffer I (NaOH 25 mM, EDTA(Na) 0.2mM, pH 12), mixed by pipetting and incubated at 94 °C for 30 min. Next, 20 μL of Buffer II (Tris-HCl 40 mM, pH 7.46) were added to the mixture and stored at −20 °C.

### Isolation and genome sequencing of *Chlamydomonas-associated* bacteria

Soil bacteria associated with *Cr* after co-cultivation were isolated from mesocosm cultures using a dilution-to-extinction approach (Bai *et al*., 2015; Wippel *et al*., 2021). Briefly, cultures containing *Cr* and bacteria from CAS soil washes as described above were incubated in TP or B&D media. After 7 days of co-cultivation mesocosm samples were fractionated by sequential centrifugation and sonication (**Fig. S1C**; Kim *et al*., 2014) prior to dilution. For fractionation, cultures were centrifuged at 400×g for 5 min to recover the supernatant. The pellet was washed with 1x TE buffer followed by sonication in a water bath at room temperature for 10 min and centrifugation at 1,000×g for 5 min. The supernatant from the first and second centrifugation were pooled together and diluted at either 1:10,000 or 1:50,000. Diluted supernatants were then distributed into 96-well microtiter plates containing 20% TSB media. After 3 weeks of incubation in the dark at room temperature, plates that showed visible bacterial growth were chosen for *16S* rRNA amplicon sequencing. For identification of the bacterial isolates, a two-step barcoded PCR protocol was used as previously described (Wippel *et al*., 2021). Briefly, DNA extracted from the isolates was used to amplify the v5-v7 fragments of the *16S* rRNA gene by PCR using the primers 799F (AACMGGATTAGATACCCKG) and 1192R (ACGTCATCCCCACCTTCC), followed by indexing of the PCR products using Illumina-barcoded primers. The indexed *16S* rRNA amplicons were subsequently pooled, purified, and sequenced on the Illumina MiSeq platform. Next, cross-referencing of IPL sequences with mesocosm profiles allowed us to identify candidate strains for further characterization, purification, and whole-genome sequencing. Two main criteria were used for this selection: first, we aimed at obtaining maximum taxonomic coverage and selected candidates from as many taxa as possible; second, we gave priority to strains whose *16S* sequences were highly abundant in the natural communities. Whenever multiple candidates from the same phylogroup were identified, we aimed at obtaining multiple independent strains, if possible, coming from separate biological replicates to ensure they represented independent isolation events. After validation of selected strains, 185 were successfully subjected to whole-genome sequencing. Liquid cultures or swabs from agar plates from selected bacterial strains (**Supplementary Data 3**) were used to extract DNA using the QiAmp Micro DNA kit (Qiagen, Hilden, Germany). The extracted DNA was treated with RNase, and purified. Quality control, library preparation, and sequencing (2 x 150 bp; Illumina HiSeq3000) at a 4-5 million reads per sample were performed by the Max Planck-Genome Center, Cologne, Germany (https://mpgc.mpipz.mpg.de/home/).

### Multi-species microbiota reconstitution experiments

The gnotobiotic FlowPot (Kremer *et al*., 2021) system was used to grow *Cr* or *A. thaliana* plants with and without bacterial SynComs. This system allows for even inoculation of each FlowPot with microbes by flushing of the pots with the help of a syringe attached to the bottom opening. After FlowPot assemblage, sterilization and microbial inoculation sterilized seeds were placed on the matrix (peat and vermiculite, 2:1 ratio), and pots were incubated under short-day conditions (10 hours light, 21°C; 14 hours dark, 19°C), standing in customized plastic racks in sterile ‘TP1600+TPD1200’ plastic boxes with filter lids (SacO2, Deinze, Belgium). For SynCom preparation, bacterial strains from either *Cr*- or *At*-SPHERE were grown separately in liquid culture for 2-5 days in 50% TSB media and then centrifuged at 4,000 xg for 10 min and re-suspended in 10 mM MgCl_2_ to remove residual media and bacteria-derived metabolites. Equivalent ratios of each strain, determined by optical density (OD600) were combined to yield the desired SynComs (**Table S1**). An aliquot of the SynComs as reference samples for the experiment microbial inputs were stored at −80°C for further processing. SynCom bacterial cells (10^7^) were added to either 50 mL of TP10 or ½ MS (Duchefa Biochemie, Haarlem, Netherlands), which were then inoculated into the FlowPots using a 60 mL syringe. For *Cr*-inoculated pots, 10^5^ of washed *Cr* cells were added to the 50mL of media with or without microbes to be inoculated into the FlowPots.

*Chlamydomonas* or *Arabidopsis* FlowPots were grown side-by-side in gnotobiotic boxes, with six pots in total per box. This experiment was repeated in three independent biological replicates. After five weeks of growth, roots were harvested and cleaned thoroughly from attached soil using sterile water and forceps. Surface of *Chlamydomonas* pots were used as phycosphere samples (cells were harvested from visibly green surface areas, top soil samples). In addition, to remove any possible background effect from carry-over soil particles, the surface-harvested samples were washed in sterile TE supplemented with 0.1% of Triton X-100 by manually shaking in 2-mL Eppendorf tubes. Then, the tubes rested for a few minutes and the supernatant was used as “cell fraction” samples. Finally, soil from unplanted pots were collected as soil samples and treated similarly as *Chlamydomonas*-inoculated pots for microbial community comparison. All phycosphere, root (comprising both the epiphytic, and endophytic compartments), and soil (soil from unplanted pots) samples were transferred to Lysing Matrix E tubes (MP Biomedicals, Solon, USA), frozen in liquid nitrogen, and stored at −80°C for further processing. DNA was isolated from those samples using the MP Biomedicals FastDNA™ Spin Kit for Soil, and from the input SynCom by alkaline lysis, and subjected to bacterial community profiling.

To ensure sufficient surface for phycosphere harvesting, we set up an additional experiment based on sterile peat without FlowPots. Experiments with the mixed SynCom of *Cr*- and *At*-SPHERE strains were conducted using sterile ‘TP750+TPD750’ plastic boxes (SacO2, Deinze, Belgium). Sterile soil and vermiculite were mixed in a 2:1 ratio and added to each box. Next, the boxes were inoculated by adding 95 mL of TP10 or ½ MS, for the *Chlamydomonas* or *Arabidopsis* boxes respectively, containing 2×10^7^ bacterial cells.

Samples for chlorophyll extraction were collected from the different *Chlamydomonas* containing gnotobiotic systems by harvesting the green surface of the peat and extracting the cells as described above. Then, 1 mL of these extracts were centrifuged at 14,000 xg for 1 min at 4°C with 2.5 μl 2% (v/v) Tween 20 (Sigma-Aldrich, Darmstadt, Germany) to promote the aggregation into a pellet. Then, the supernatant was completely removed and the pellets stored at −80°C until extraction.

### Chlorophyll extraction from algae-containing samples

From each extracted cell samples from the gnotobiotic soil system, 1 mL was collected and mixed with 2.5 μL of 2% (v/v) Tween 20 in 1.5 mL Eppendorf tubes. The samples were centrifuged for 1 min at 14,000×g and 4 °C, then the supernatant was removed and the pellet stored at −80 °C. Frozen samples were thawed on ice for 2 min and 1 mL of HPLC grade methanol (Sigma, 34860-4L-R) added to the pellets. The tubes were covered from the light using aluminum foil and mixed using the vortex for 1 min. After vortexing, the cells were incubated in the dark at 4 °C for five minutes. Next, the pigments were obtained by centrifuging the cells for 5 minutes at maximum speed and 4 °C and recovering the supernatant. The pigments absorbance at 652 and 665 nm was measured in a plate reader Infinite M200Pro using methanol as blank. The absorbance values were then substituted in the following equation Chl a + Chlb = 22.12×Abs652 + 2.71×Abs665 (Porra *et al*., 1989).

### Split co-cultivation system

Co-cultivation devices were built by adapting 150 mL Stericup-GV filtration devices (Merck Millipore, Darmstadt, Germany) harboring a 0.22 μm filter membrane (Alvarez and Cava, 2018). Each co-cultivation device was assembled inside a clean hood 150 mL and 100 mL of TP10 were added into the big and small chamber of the filtration device, respectively. Chambers were inoculated at different cell concentrations depending on the content of the chamber (**Fig S1F**). The concentrations used were 10^5^ and 10^7^ cells/mL for *Chlamydomonas* and SynCom respectively. For the C+SC condition, the inoculum concentration was the same as for individual content chambers. After inoculation the devices were transferred to a shaking platform and incubated under the same conditions used for *Cr* liquid cultures described above. Four samples per chamber were harvested for DNA extraction, fluorescence, and cell growth at the start of the incubation and 7 days after inoculation. These experiments were repeated in three independent biological replicates, containing one technical replicate each.

Additionally, a full-factorial replicate of the experiment was carried out using a custom-made co-cultivation device (Cat. #0250 045 25, WLB Laborbedarf, Möckmühl, Germany). Briefly, two 250 mL borosilicate glass bottles (**Fig. S1F**) were modified by adding on the sidewall of each bottle a glass neck with a NW25 flange. The flange holds a disposable 0.22 μm-pore PVDF Durapore filtration membrane (Merck Millipore, Darmstadt, Germany) and is kept in place by an adjustable metal clamp. In this device, each bottle holds 150 mL of TP10 and the initial cell concentrations were the same as the ones used in the previously described co-cultivation device. Similar to the Stericup system, four samples per chamber were harvested for DNA extraction. Chlorophyll fluorescence and cell growth measurements were collected at the start of the incubation and 7 days after inoculation. These experiments were repeated in three independent biological replicates, containing one technical replicate each.

### Preparation of SynCom inocula

Bacterial cultures from the strains selected for the different SynComs (**Supplementary Data 4**) were started from glycerol stocks which were used to streak agar plates containing TSA 50% media. Plates were cultured at 25 °C for five days and later used to inoculate culture tubes with 1 mL of 50% TSB media. The tubes were incubated for six days at 25 °C and 180 RPM. After 6 days, the cultures were washed three times by centrifugation at 4,000 ×g for 5 min, the supernatant discarded, and the pellet resuspended into 2 mL of TP or TP10 media. The washed cultures were further incubated with shaking at 25 °C for an additional day. Bacterial concentration in washed cultures was determined by measuring OD_600_ and, subsequently pooled in equal ratios. Cell counts of the pooled SynCom were measured using the Multisizer 4e and adjusted to 10^6^, to inoculate together with 10^4^ cells of *Cr* (prepared as described above) in 50 mL of TP10 in 200-mL flasks. These flasks were inoculated in triplicate and three biological replicates were prepared for both bacteria and *Cr* start inocula. As controls, *Cr*-only cultures and SynCom-only cultures were incubated in parallel, and samples taken at 0, 1, 4, 7 for community profiling, and at 0, 4, 7, 14 days for *Cr* cell counts.

### Culture-independent bacterial *16S* rRNA sequencing

DNA samples were used in a two-step PCR amplification protocol. In the first step, V2–V4 (341F: CCTACGGGNGGCWGCAG; 806R: GGACTACHVGGGTWTCTAAT) or V4-V7 (799F: AACMGGATTAGATACCCKG; 1192R: ACGTCATCCCCACCTTCC) of bacterial *16S* rRNA were amplified. Under a sterile hood, each sample was amplified in triplicate in a 25 μL reaction volume containing 2 U DFS-Taq DNA polymerase, 1x incomplete buffer (Bioron GmbH, Ludwigshafen, Germany), 2 mM MgCl2, 0.3% BSA, 0.2 mM dNTPs (Life technologies GmbH, Darmstadt, Germany) and 0.3 μM forward and reverse primers. PCR was performed using the same parameters for all primer pairs (94°C/2 min, 94°C/30 s, 55°C/30 s, 72°C/30 s, 72°C/10 min for 25 cycles). Afterwards, single-stranded DNA and proteins were digested by adding 1 μL of Antarctic phosphatase, 1 μL Exonuclease I and 2.44 μL Antarctic Phosphatase buffer (New England BioLabs GmbH, Frankfurt, Germany) to 20 μl of the pooled PCR product. Samples were incubated at 37°C for 30 min and enzymes were deactivated at 85°C for 15 min. Samples were centrifuged for 10 min at 4,000 rpm and 3 μL of this reaction were used for a second PCR, prepared in the same way as described above using the same protocol but with cycles reduced to 10 and with primers including barcodes and Illumina adaptors. PCR quality was controlled by loading 5 μL of each reaction on a 1% agarose gel and affirming that no band was detected within the negative control. Afterwards, the replicated reactions were combined and purified. In the case of bacterial amplicons with possible plant DNA PCR products, amplicons were loaded on a 1.5% agarose gel and run for 2 hours at 80 V. Subsequently, bands with a size of ~500 bp were cut out and purified using the QIAquick gel extraction kit (Qiagen, Hilden, Germany). If plant DNA PCR products were not present, bacterial amplicons were purified with Agencourt AMPure XP beads DNA concentration was fluorescently determined, and 30 ng DNA of each of the barcoded amplicons were pooled in one library. The library was then purified and re-concentrated twice with Agencourt AMPure XP beads, and pooled in similar ratios for sequencing. Paired-end Illumina sequencing was performed in-house using the MiSeq sequencer and custom sequencing primers (**Supplementary Data 5**).

### Analysis of culture-independent bacterial *16S* rRNA profiling

Amplicon sequencing data from *Cr* or *At* roots grown in CAS soil in the greenhouse, along with unplanted controls, were demultiplexed according to their barcode sequence using the QIIME pipeline (Caporaso *et al*., 2010). Afterwards, DADA2 (Callahan *et al*., 2016) was used to process the raw sequencing reads of each sample. Unique amplicon sequencing variants (ASVs) were inferred from error-corrected reads, followed by chimera filtering, also using the DADA2 pipeline. Next, ASVs were aligned to the SILVA database (Quast *et al*., 2013) for taxonomic assignment using the naïve Bayesian classifier implemented by DADA2. Raw reads were mapped to the inferred ASVs to generate an abundance table, which was subsequently employed for analyses of diversity and differential abundance using the R package *vegan* (Oksanen *et al*., 2019).

Amplicon sequencing reads from the *Cr* IPL and from the corresponding mesocosm culture-independent community profiling were quality-filtered and demultiplexed according to their two-barcode (well and plate) identifiers using custom scripts and a combination of tools included in the QIIME and USEARCH (Edgar *et al*., 2010) pipelines. Next, sequences were clustered into Operational Taxonomic Units (OTUs) with a 97% sequence identity similarity using the UPARSE algorithm, followed by identification of chimeras using UCHIME (Edgar *et al*., 2011). Samples from wells with fewer than 100 good quality reads were removed from the data set as well as OTUs not found in a well with at least ten reads. Recovery rates (**Figure S4A**). were estimated by calculating the percentage of the top 100 most abundant OTUs found in natural communities (greenhouse experiment) that had at least one isolate in the culture collection (62%), and the total aggregated relative abundances of recovered OTUs (63%). We identified IPL samples matching OTUs found in the culture-independent root samples and selected a set of 185 representative strains maximizing taxonomic coverage for subsequent validation and whole-genome sequencing, forming the basis of the *Cr*-SPHERE collection.

### Meta-analysis of phycosphere and root microbiota profiles

For the meta-analysis of root microbiota samples across plant species, data from previous studies of *Arabidopsis* and *Lotus* grown in CAS soil (Thiergart *et al*., 2019; Harbort *et al*., 2020) were processed using the pipeline described above and merged with samples obtained from the Cooloola natural site chronosequence (Yeoh *et al*., 2017). Sequencing reads from this latter study (Roche 454) were quality filtered and trimmed after removal of primer sequences. Given that these studies employed non-overlapping sequencing primers, all datasets were combined after aggregating relative abundances at the bacterial order taxonomic level. The core taxa of the root microbiota were determined by identifying bacterial orders present in every plant species with an occupancy of at least 80% (i.e., found in at least 80% of the root samples of a given species) with a relative abundance above 0.1%. To infer the phylogenetic relationship between the different hosts, protein sequences of the ribulose-bisphosphate carboxylase (*rbcL*) gene for *Cr* and the 35 analyzed plant species were recovered from GenBank. The sequences were aligned using Clustal Omega (Sievers *et al*., 2011) with default parameters, and the alignment used to infer a maximum likelihood phylogeny using FastTree (Price *et al*., 2010).

### Analysis of culture-dependent amplicon sequencing data

Sequencing data from SynCom experiments was pre-processed similarly as natural community *16S* rRNA data. Quality-filtered, merged paired-end reads were then aligned to a reference set of sequences extracted from the whole-genome assemblies of every strain included in a given SynCom, using Rbec (Zhang *et al*., 2021b). We then checked that the fraction of unmapped reads did not significantly differ between compartment, experiment or host species. Next, we generated a count table that was employed for downstream analyses of diversity with the R package vegan. Finally, we visualized amplicon data from all experimental systems using the *ggplot2* R package (Wickham *et al*., 2016).

### Genome assembly and annotation

Paired-end Illumina reads were first trimmed and quality-filtered using Trimmomatic (Bolger *et al*., 2014). QC reads were assembled using the IDBA assembler (Peng *et al*., 2012) within the A5 pipeline (Tritt *et al*., 2012). Assembly statistics and metadata from the assembled genomes can be found in **Supplementary Data 3**. Genome assemblies with either multi-modal *k*-mer and G+C content distributions or multiple cases of marker genes from diverse taxonomic groups were flagged as not originating from clonal cultures. Such assemblies were then processed using a metagenome binning approach (Pasolli *et al*., 2019). Briefly, contigs from each of these samples were clustered using METABAT2 (Kang *et al*., 2019) to obtain metagenome-assembled genomes (MAGs). Each MAG was analyzed to assess completeness and contamination using CheckM (Parks *et al*., 2015). Only bins with completeness scores larger than 75% and contamination rates lower than 5% were retained and added to the collection (**Supplementary Data 3**; designated MAG in the column ‘type’). Classification of the bacterial genomes into phylogroups was performed by calculating pair-wise average nucleotide identities using FastANI (Jain *et al*., 2018) and clustering at a 97% similarity threshold. Functional annotation of the genomes was conducted using Prokka (Seeman *et al*., 2014) with a custom database based on KEGG Orthologue (KO) groups (Kanehisa *et al*., 2014) downloaded from the KEGG FTP server in November 2019. Hits to sequences in the database were filtered using an *E*-value threshold of 10 × 10-9 and a minimum coverage of 80% of the length of the query sequence.

### Comparative genome analyses of the *Cr*-, *At*- and *Lj*-SPHERE culture collections

The genomes from the *Cr*-, *At*- and *Lj*-SPHERE culture collections (Bai *et al*., 2015; Wippel *et al*., 2021) were queried for the presence of 31 conserved, single-copy marker genes, known as AMPHORA genes (Wu *et al*., 2008). Next, sequences of each gene were aligned using Clustal Omega (Sievers *et al*., 2011) with default parameters. Using a concatenated alignment of each gene, we inferred a maximum likelihood phylogeny using FastTree (Price *et al*., 2010). This tree was visualized using the Interactive Tree of Life web tool (Letunic *et al*., 2019). Finally, genomes from the three collections (*Cr*-SPHERE, *At*-SPHERE and *Lj*-SPHERE) were clustered into phylogroups, roughly corresponding to a species designation (Olm *et al*., 2020) using FastANI (Jain *et al*., 2018) and a threshold of average nucleotide identity at the whole genome level of at least 97%. Functional comparison among the genomes from the *Cr*-, *Lj*- and *At*-SPHERE collections was performed by comparing their annotations. KO groups were gathered from the genome annotations and aggregated into a single table. Lastly, functional distances between genomes based on Pearson correlations were used for principal coordinate analysis using the *cmdscale* function in R.

## Data deposition

Raw sequencing will be deposited into the European Nucleotide Archive (ENA) under the accession number PRJEB43117. The scripts used for the computational analyses described in this study are available at http://www.github.com/garridoo/crsphere, to ensure replicability and reproducibility of these results.

## Author contributions

P.D., J.F.-U., K.W. and R.G.-O. designed the experiments. P.D. and J.F.-U. conducted the greenhouse experiments. P.D. and K.W. performed the mesocosm experiments. P.D., J.F.-U., and K.W. established the IPL bacterial library and characterized the *Cr*-SPHERE core culture collection. P.D. and J.F.-U. performed the synthetic community experiments. P.Z., J.F.-U. and R.G.-O. analyzed whole-genome sequencing data. P.D., J.F.-U., P.Z., R.G. and R.G.-O. analyzed bacterial *16S* rRNA amplicon data. P.D., J.F.-U., K.W. and R.G.-O interpreted the results. P.D., J.F.-U. and R.G.-O wrote the paper.

## Acknowledgements

We would like to thank Dr. Paul Schulze-Lefert for his support and advice throughout the duration of this project. We would also like to thank Dr. Michael Melkonian, Dr. Stephane Hacquard, Dr. Thomas Nakano, Dr. Oliver Ebenhöh, and Dr. Andreas Weber for their critical comments on this manuscript. We thank Dr. Ralph Bock, Dr. Juliane Neupert and Dr. Ru Zhang for their advice and assistance during the early stages of this research project. Finally, we thank Rozina Kardarakis for grammatical and style corrections of the manuscript. This research was funded by the Max Planck Society and the Deutsche Forschungsgemeinschaft (DFG, German Research Foundation) under Germany’s Excellence Strategy – EXC-Nummer 2048/1–project 390686111, and the ‘2125 DECRyPT’ Priority Programme.

## Notes

### Competing Interest Statement

The authors have declared no competing interest.

