## Supplementary Information for "Characterization of the *Chlamydomonas reinhardtii* phycosphere reveals conserved features of the plant microbiota"

### **Supplementary Figures**

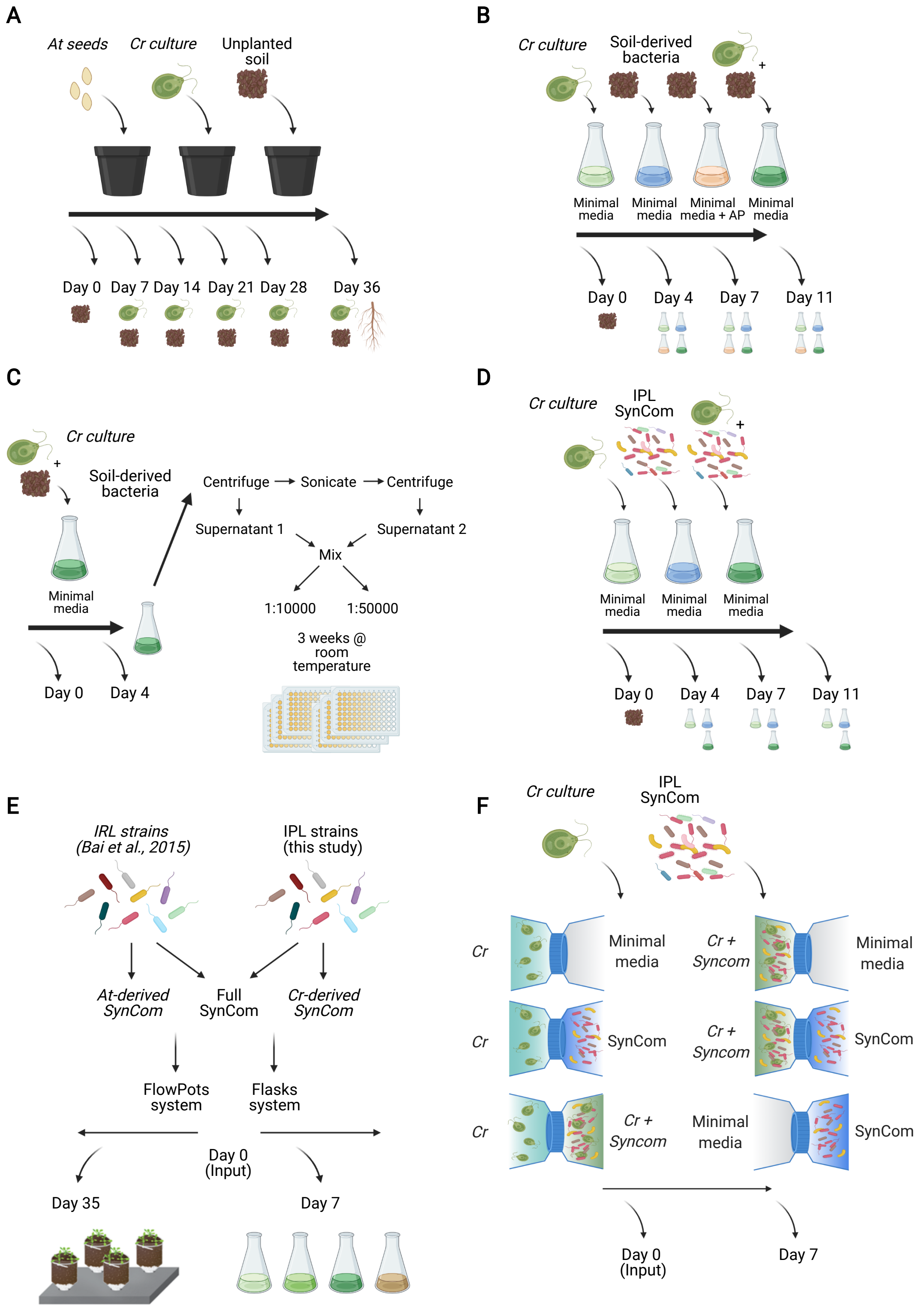


**Figure S1. Schematic description of the experimental approaches employed in this study (caption on next page).**

**Figure S1. Schematic description of the experimental approaches employed in this study (figure on previous page).**

(**A**) Greenhouse experiment. Pots containing CAS natural soil were either sown with *At* seeds, inoculated with *Cr* cultures or mock-treated. Samples were taken over time for bacterial community profiling. (**B**) Mesocosm experiments. In different liquid media (minimal media or AP-containing media), *Cr* cultures were co-incubated with soil-derived bacteria. Samples were taken over time for bacterial community profiling and assessment of *Cr* growth. (**C**) Protocol for the establishment of the *Cr*-IPL bacterial culture collection generation. *Cr* cultures were co-incubated with soil-derived bacteria for 7 days and fractionated to enrich for phycosphere bacteria. This phycosphere fraction was then diluted and incubated for several weeks in 96-well plates. Subsequently, bacterial cultures were subjected to *16S* rRNA amplicon profiling for further analysis (**D**) SynCom reconstitution experiment. From the core culture *Cr*-SPHERE collection, 26 representative strains were selected, pooled together and co-incubated with *Cr* in minimal media for 7 days. (**E**) Cross-inoculation experiment. One bacterial strain per shared bacterial family from the *At*-SPHERE and *Cr*-SPHERE were selected and assembled into root or phycosphere SynComs. These SynComs were cross-inoculated in *At* and *Cr* in two gnotobiotic systems (i.e. soil-based FlowPots and liquid-based flasks), individually or in a mixed community. Root and phycosphere samples were harvested after 5 weeks of co-incubation for community profiling and assessment of *Cr* growth. (**F**) Split co-cultivation system. Co-cultivation chamber separated by a 0.22 um filter were used to inoculate *Cr*, the phycosphere SynCom alone or both together in different combinations. Samples were harvested after 7 days for community profiling and *Cr* growth measurements.

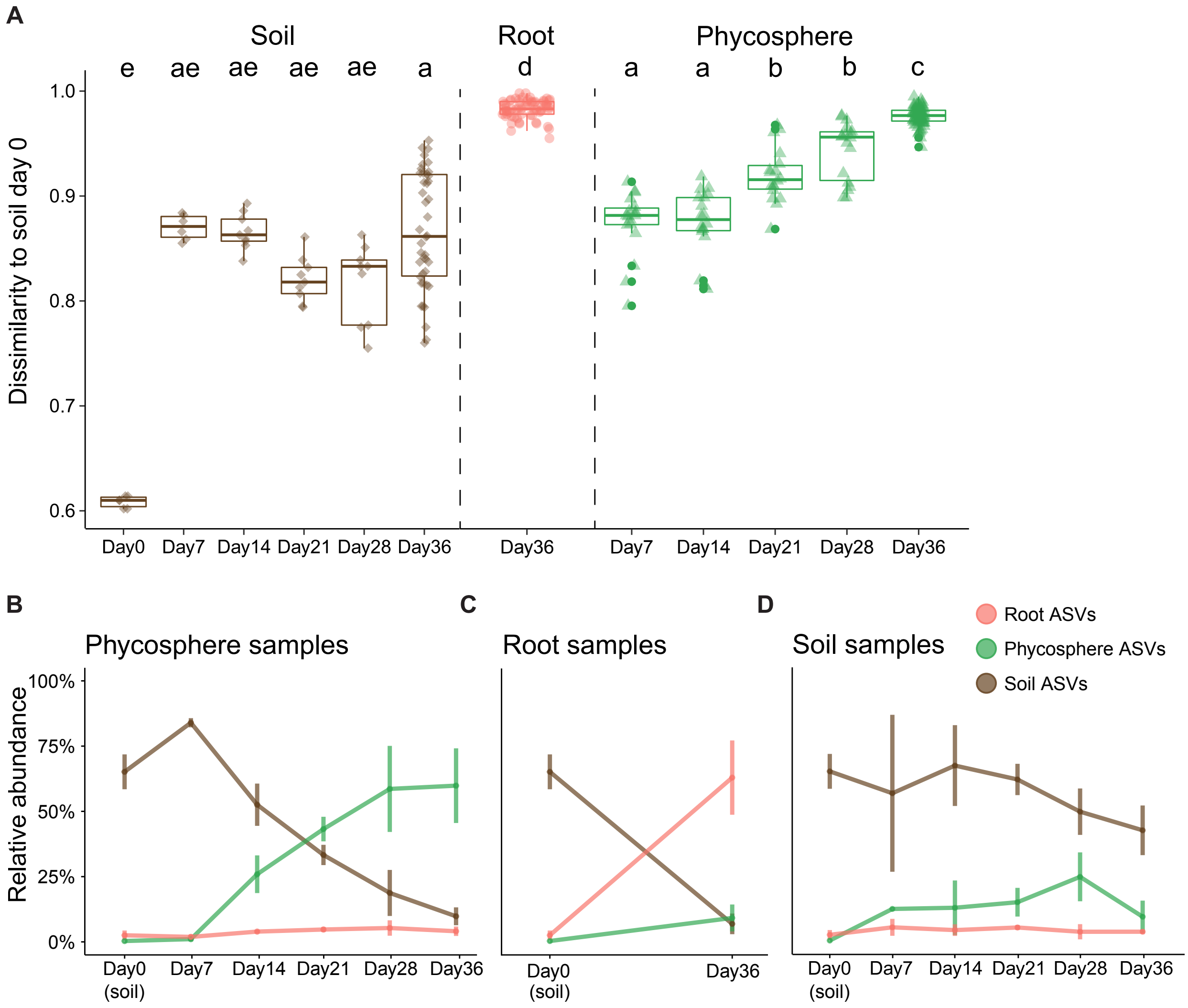


**Figure S2. Culture-independent analysis of phycosphere- and root-associated communities in a natural soil.**

(**A**) Bray-Curtis dissimilarities between phycosphere, root and soil communities, compared to the initial soil input (day 0). Boxplots are color-coded depending on the fraction. Significant differences are marked with different letters (Kruskal-Wallis, followed by a Dunn's *post hoc*, with Bonferroni correction). (**B-D**) Dynamics of relative abundances of ASVs enriched in phycosphere (**B**), root (**C**) or soil samples (**D**) over time, compared to initial soil input (day 0; Wilcoxon test; $P<0.05$). Curves are color-coded depending on the fraction indicated.

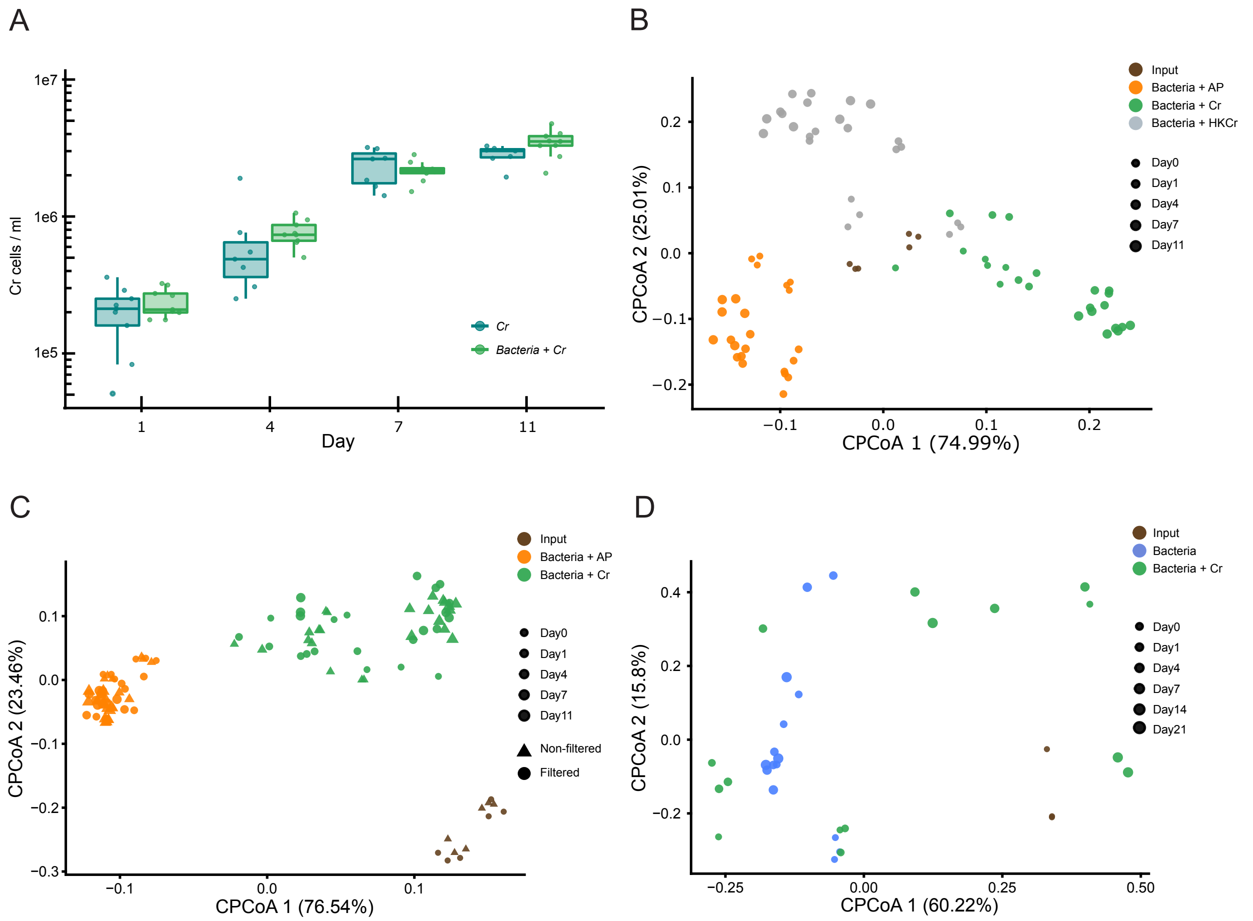


**Figure S3. Analysis of phycosphere communities in mesocosm experiments.**

(**A**) Cell counts of axenic *Cr* cultures (dark green) and of *Cr* co-inoculated with soil-derived bacteria (light green) over time. No significant differences were found between growth conditions at each time point (Wilcoxon test). (**B**) PCoA of Bray-Curtis dissimilarities constrained by the experimental condition (17.5% of variance; $P$<0.001) of soil-derived bacterial communities co-inoculated with AP, live *Cr* cultures or heat-killed *Cr*. Points in the plot are color-coded according to the condition and their size indicates the time point at which samples were harvested. (**C**) PCoA of Bray-Curtis dissimilarities constrained by the experimental condition (17.5% of variance; $P$<0.001) of filtered (circles) or non-filtered (triangles) bacterial communities derived from soil and co-inoculated with AP or *Cr* cultures. No significant separation of samples was found based on filtration treatment (0.73% of variance). (**D**) PCoA of Bray-Curtis dissimilarities constrained by experimental condition and time point of (40.1% of variance; $P$<0.005) of soil-derived bacterial communities either co-inoculated with *Cr* cultures or alone in minimal media, under day/night light conditions.

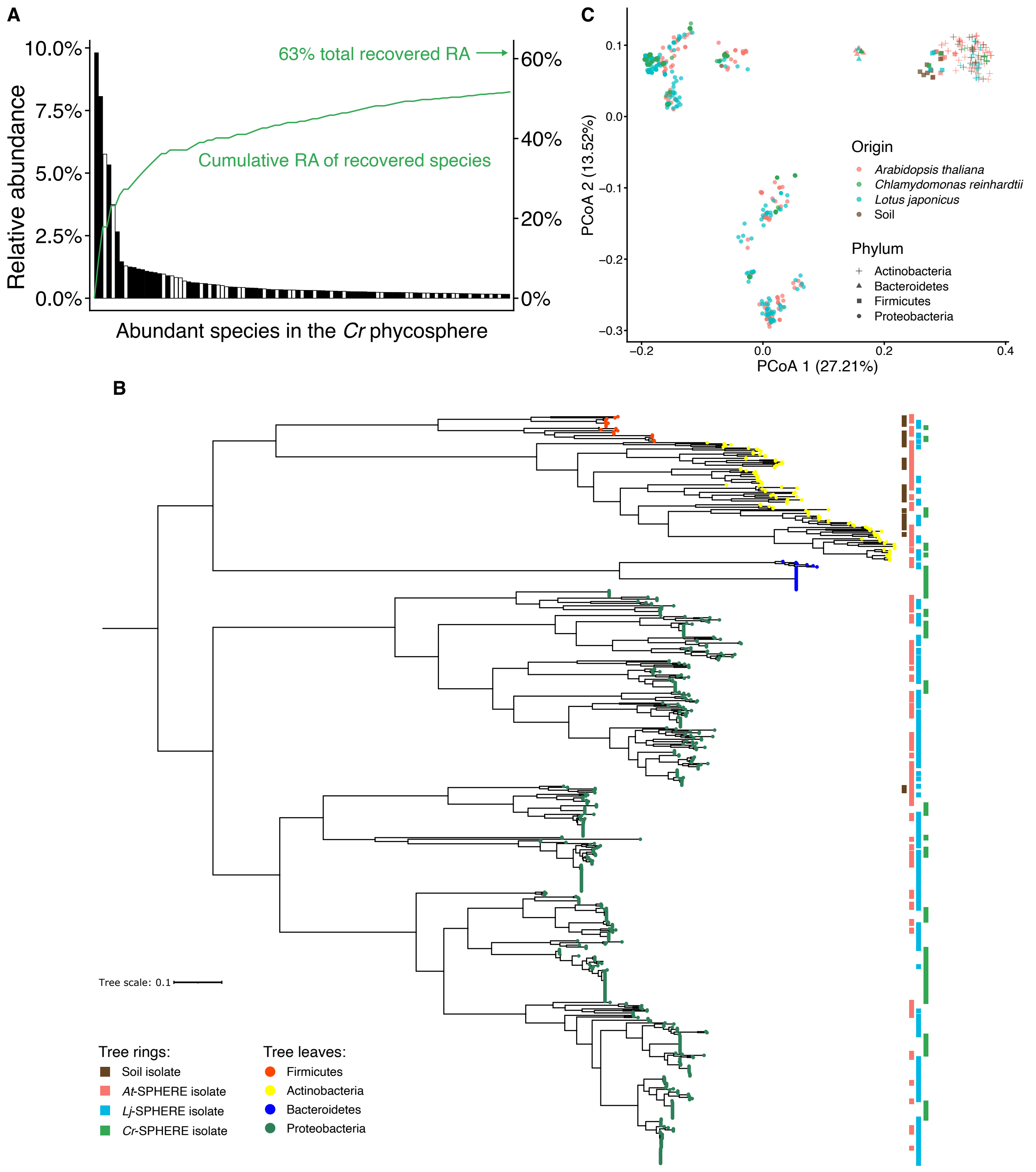


**Figure S4. Characterization of the IPL and *Cr*-SPHERE culture collection (caption on next page).**

**Figure S4. Characterization of the IPL and *Cr*-SPHERE culture collection (figure on previous page).**

(**A**) Bar plot showing the relative abundance of the top 100 most abundant OTUs found in culture-independent phycospheres (greenhouse experiment) indicating whether a representative exemplar was recovered in the IPL bacterial library (62%). The cumulative relative abundance curve and arrow represent total contribution of recovered OTUs to the culture-independent phycosphere communities (63% of the entire community). (**B**) Whole-genome phylogenetic tree of bacterial strains derived from soil or the *At*-, *Lj*-, *Cr*-SPHERE culture collections based on a concatenation of single copy phylogenetic markers known as AMPHORA genes. Tree leaves are colored by taxonomic affiliation, while colored boxes represent the origin of each genome. (**C**) PCoA of functional distances of genomes from bacterial isolates of the *Chlamydomonas* (Cr-SPHERE), *Arabidopsis* (At-SPHERE), and *Lotus* (Lj-SPHERE) bacterial culture collections.

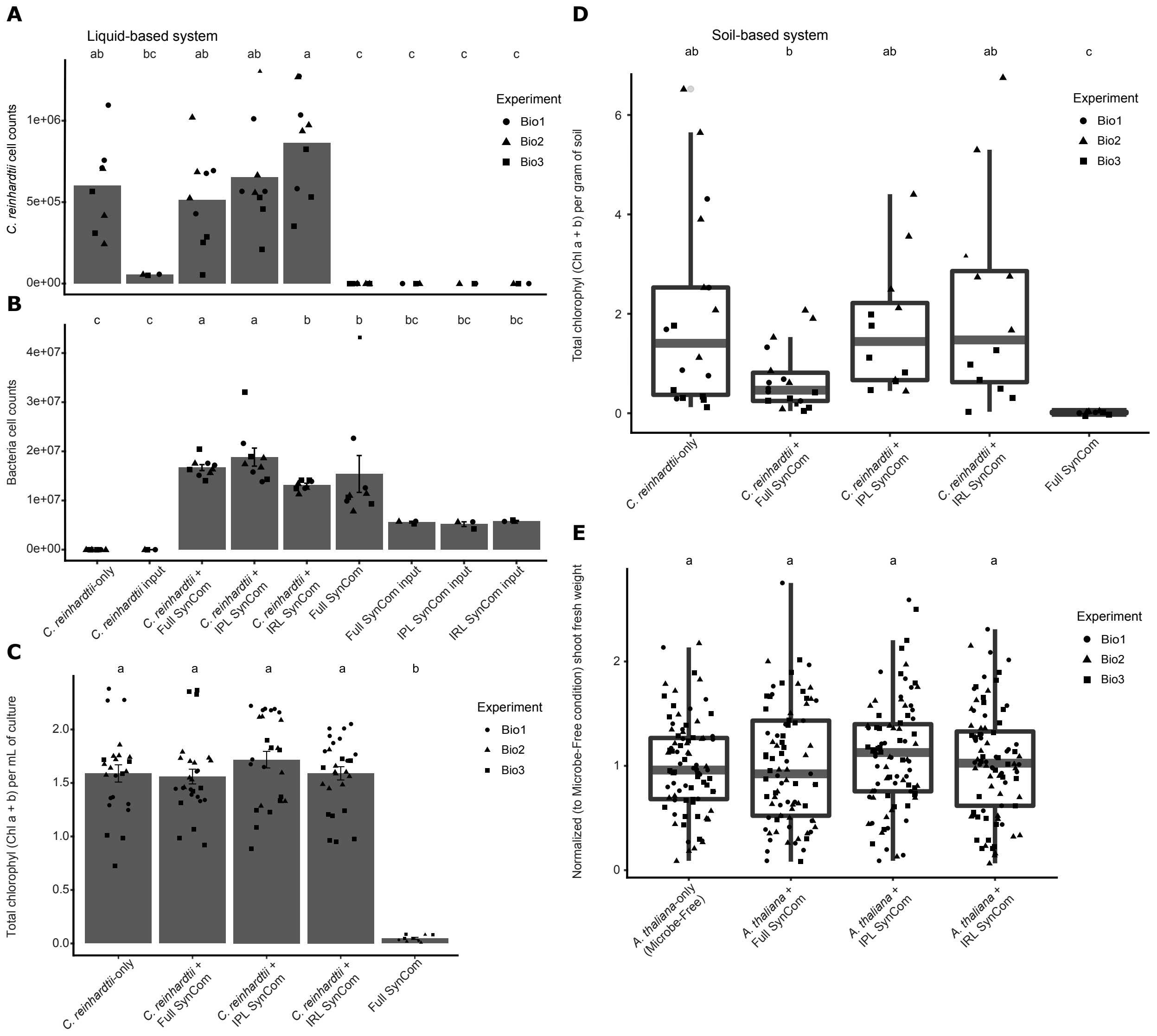


**Figure S5. Bacterial and host growth parameters in cross-inoculation experiments with IPL and IRL bacterial SynComs.**

(**A**) *Cr* cell counts after 7 days of co-incubation with different bacterial SynCom combinations in liquid system. (**B**) Cell counts of different bacterial SynCom combinations after 7 days of co-incubation with or without *Cr* in a liquid system. (**C**) *Cr* total chlorophyll content after 7 days of co-incubation with different bacterial SynCom combinations in the liquid system. (**D**) Cr total chlorophyll content after 4 weeks of co-incubation with different bacterial SynCom combinations in the soil-based system. (**E**) *At* normalized shoot fresh wait after 4 weeks of co-incubation with different bacterial SynCom combinations in the soil-based system. Letters on top indicate significant differences between condition within one culture collection (Kruskal-Wallis, followed by a Dunn's test *post hoc*, and Bonferroni correction).

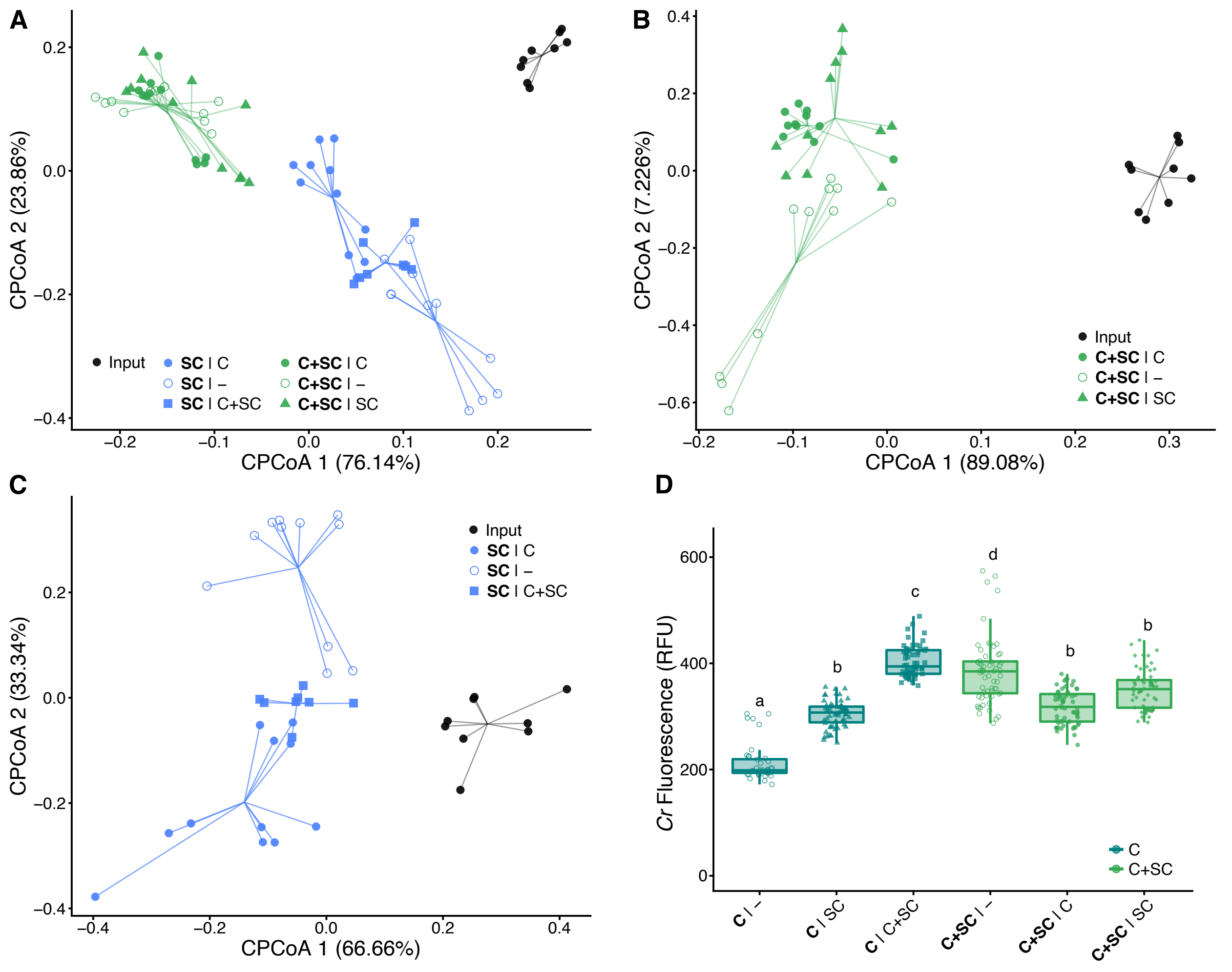


**Figure S6. Effect of physical proximity in SynCom community structure and *Cr* growth (split bottle system).**

(**A**) PCoA plot of Bray-Curtis dissimilarities constrained by physical proximity to *Cr* (20.6% variance; $P<0.001$). Points are color-coded based on the community present in a given compartment, and shapes depict the content of the neighboring compartment. (**B-C**) PCoA plot of Bray-Curtis distances constrained by physical proximity to *Cr* of C+SC (**B**; 39.7% variance; $P<0.001$) and SC-only compartments (**C**; 22.7% variance; $P<0.001$). (**D**) Cr growth after 7 days of co-incubation with different compartment combinations, measured as relative chlorophyll fluorescence (RFU). Boxplots are color-coded depending on whether *Cr* was incubated alone in a given compartment (dark green) or in combination with SC (light green).
